# Non-apoptotic caspase activation sustains ovarian somatic stem cell functions by modulating Hedgehog-signalling and autophagy

**DOI:** 10.1101/722330

**Authors:** Alessia Galasso, Daria Iakovleva, Luis Alberto Baena-Lopez

## Abstract

There is increasing evidence associating the activity of caspases with the regulation of basic cellular functions beyond apoptosis. Accordingly, the dysregulation of these novel non-apoptotic functions often sits at the origin of neurological disorders, metabolic defects, autoimmunity, and cancer. However, the molecular interplay between caspases and the signalling networks active in non-apoptotic cellular scenarios remains largely unknown. Our work show that non-apoptotic caspase activation is critical to modulate Hedgehog-signalling and autophagy in ovarian somatic cells from both *Drosophila* and humans under moderate stress. We also demonstrate that these novel caspase functions are key to sustain stem cell proliferation and differentiation without inducing apoptosis. Finally, we molecularly link these caspase-dependent effects to the fine-tuning of the Hedgehog-receptor, Patched. Together, these findings confer a pro-survival role to the caspases, as opposed to the widely held apoptotic function assigned to these enzymes.

## INTRODUCTION

In recent years, expanding evidence is indicating the ability of caspases to regulate essential cellular functions beyond apoptosis^1–3^. Accordingly, these novel non-apoptotic caspase roles ensure tissue homeostasis, whilst preventing diseases^1,2^. However, the molecular characterisation of these alternative caspase functions remains largely unknown in the vast majority of cell types, including stem cells. During the last decade, the investigations using the adult *Drosophila* ovary have illuminated fundamental principles of stem cell physiology and intercellular communication^4^. Interestingly, these progenitor cells and their progeny can activate caspases at sublethal levels in response to robust environmental stress^5^. Therefore, it is an ideal cellular system to study the interplay between caspases, signalling mechanisms, and stem cell physiology.

The early development of *Drosophila* female gametes occurs in a cellular structure referred to as the germarium. The germarium is formed by the germline and the surrounding somatic cells^4^ (Fig. 1a). The cellular properties within the germarium are strongly defined by the Hedgehog-signalling pathway^6–11^. The interaction of the Hedgehog (Hh) ligand with its membrane receptor Patched (Ptc) allows the activation of the signalling transducer Smoothened (Smo)^12^. This prevents the proteolytic processing of the transcriptional regulator Cubitus interruptus (Ci)^12^, thus eliciting the activation of tissue-specific target genes^12^. The main somatic Hh-receiving cells in the germarium are the escort cells^9^ and the follicular stem cells^10^ (Fig. 1a). As opposed to Hh-signalling deprivation^6,8,10^, the overactivation of Hh-pathway facilitates cell proliferation and cell differentiation in the follicular stem cells and their progeny^8,11,13,14^. Beyond the developmental requirements, Hh-signalling also prevents the excess of Ptc-induced autophagy under stress conditions, thus ensuring the homeostasis of ovarian somatic cells^14,15^. Importantly, the pro-proliferative and differentiating roles of Hh-pathway are largely conserved in human ovarian cells with somatic origin^16–19^.

**Fig. 1.**
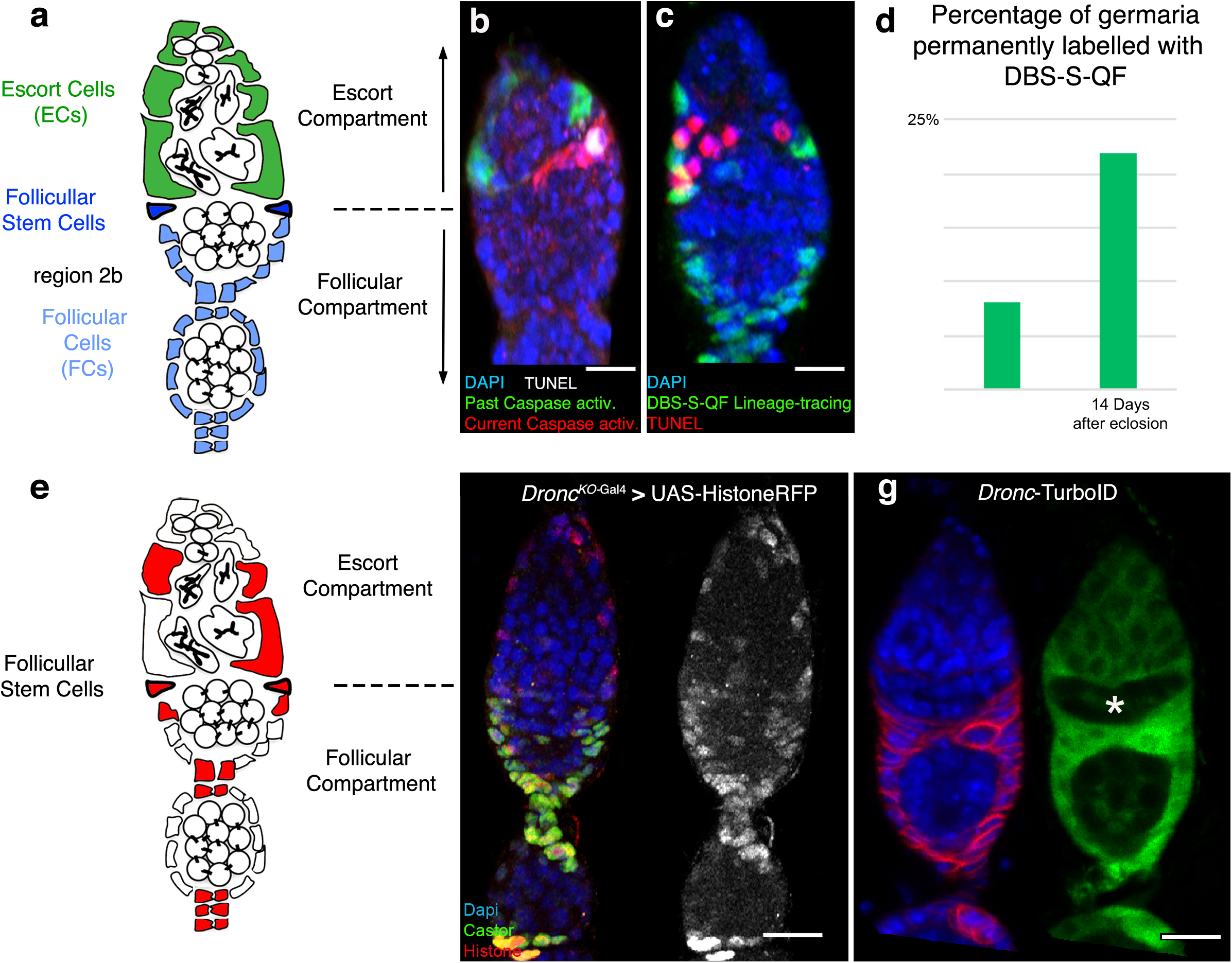
Non-apoptotic activation of initiator caspases in somatic cells of the *Drosophila* germarium. **a**. Schematic drawing of the Drosophila germarium. Somatic cells relevant for this study (escort, follicular stem and follicular) are respectively depicted in green, dark blue, light blue; germline cells are shown in white. **b**. Representative confocal image showing past (green channel, arrows) and present (red channel) caspase activation in presumptive escort somatic cells using the DBS-S-QF sensor; TUNEL staining indicates apoptosis (gray, arrowhead); Dapi labels the DNA in the entire Figure. Scale bars represents 10 μm in the entire Figure. Experimental flies were kept for 14 days at 29°C after eclosion and prior dissection. **c**. Representative confocal image showing escort and follicular somatic cells permanently labelled with DBS-S-QF sensor (green channel, arrows); the arrowhead indicates the presence of apoptotic germline cells (rey channel, TUNEL staining, arrowhead). Notice the lack of TUNEL signal in somatic cells labelled with DBS-S-QF sensor (green). Experimental flies were kept for 14 days at 29°C after eclosion and prior dissection. **d**. Graph bar indicating the percentage of ovarioles permanently labelled with DBS-S-QF sensor at 7 and 14 days; flies were raised at 18°C until eclosion, then shifted to 29°C until the indicated dissection times. **e**. Diagram depicting the presumptive cells that transcribe *Dronc* at 29°C in the germarium (red). **f**. Representative confocal image showing escort and follicular somatic cells in the germarium expressing Histone-RFP (red channel, arrows) under the regulation of *Dronc*^*KO*-Gal4^ after 7 days at 29 °C; the follicular maker Castor is shown in green. **g**. Biotinylation signal (green) generated in the germarium by a *Dronc*-TurboID allele; notice the signal enrichment in follicular stem cells and their progeny (white arrows) as well as the relative low levels in the germline (symbols). FasIII staining (red) labels the somatic cells and Dapi labels the DNA (blue). Experimental flies were kept for 7 days at 29°C after eclosion and prior dissection.

In this manuscript, we establish that the cellular properties of ovarian somatic cells under moderate stress conditions are strongly influenced by the non-apoptotic caspase-dependent regulation of Hh-signalling and autophagy. At the molecular level, we connect this novel caspase functions with the fine-tuning of the Hh receptor, Ptc. We also provide preliminary evidence suggesting that our findings from *Drosophila* could be highly relevant in human cells with a comparable origin. Together, our observations uncover unknown features of stem cell regulation and caspase biology, whilst conferring a pro-survival role to these formerly pro-apoptotic enzymes.

## RESULTS

### There is non-apoptotic activation of *Dronc* in ovarian somatic stem cells

We recently generated a novel caspase sensor based on a cleavable, but catalytically inactive version of the effector caspase, *Drice* (Drice based sensor QF; DBS-S-QF)^20^. Amongst other applications, our reporter can provide a historical perspective of initiator caspase activation in complex *Drosophila* tissues by inducing the expression of multiple fluorescent markers with variable durability^20^ (Supplementary Fig. 1a). Since strong environmental stress (starvation and cold shock) can induce widespread non-apoptotic activation of effector caspases in the *Drosophila* ovary^5^, we sought to investigate in detail with our sensor the features of such caspase activation patterns under moderate stress. The detailed inspection of adult flies kept at 29°C confirmed the presence of initiator caspase activation in subsets of somatic cells of the germarium (red and GFP signals, Fig. 1b). Intriguingly, we noticed that subsets of escort and follicular cells could show the fluorescent signature linked to caspase activation in the past (sensor-labelled cells in green with GFP), without signs of ongoing caspase activity (sensor-labelled cells in red with Tomato-HA) or apoptosis (e.g. reporter-positive cells but TUNEL negative; Fig.1b). Confirming the healthy and even proliferative status of sensor-labelled cells, we also recovered large groups of escort and follicular cells permanently decorated with the durable cellular marker induced by DBS-S-QF (lacZ positive cells, Fig. 1c). Furthermore, the number of enduringly marked germaria with this permanent caspase-labelling system increased from 8% at 7 days to 22% at 14 days after adult eclosion, respectively (Fig. 1d). These results show for the first time the presence of transient and non-apoptotic activation of initiator caspases under moderate stress in somatic cells of the germarium, including the proliferative stem cell precursors that give rise to escort and follicular cells.

Since DBS-S-QF was specifically designed to report on the activity of initiator caspases (mainly *Dronc*, the *Drosophila* orthologue of the mammalian caspase-2/9), we sought to investigate the transcriptional regulation of *Dronc* in the ovary using a *Dronc*^KO-Gal4^ line that recapitulates its physiological expression^21^. Interestingly, *Dronc*^KO-Gal4^ induced the expression of a neutral cellular marker (Histone-RFP) in variable subsets of escort and follicular cells in the germarium (Figs. 1e, 1f), as well as the polar and stalk cells (Figs. 1f, and Supplementary Fig. 1b). Subsequent cell lineage-tracing experiments confirmed this pattern of expression (Supplementary Fig. 1c). Next, we assessed whether this transcriptional regulation led to accumulating Dronc as a protein. To that end, we used a *Dronc* allele endogenously tagged with the biotin ligase TurboID^22^. The TurboID allows the detection of low protein level concentrations through the biotinylation of peptides in close proximity to the TurboID-tagged protein^22^. In line with our previous data, we observed a *Dronc*-mediated biotinylation enrichment in the follicular cells of the germarium (Fig. 1g), stalk cells and polar cells (Supplementary Fig. 1d). Confirming the specificity of the TurboID labelling, a version of Drice fused to the TurboID^22^ labelled the follicular region but the stalk cells, and polar cells were not marked (Supplementary Fig. 1e). In addition, the signal was noticeably weaker in the germline (Supplementary Fig. 1e). Together, our results suggested that there is enriched expression and transient non-apoptotic activation of Dronc in follicular stem cells of the germarium and their progeny under moderate stress conditions.

### *Dronc* acts as a pro-survival factor that sustains follicular stem cell functions

To determine the biological significance of Dronc activation in the germarium, we generated morphogenetic mosaics using a *Dronc*^l29^ null allele. These genetic mosaics caused morphological alterations and differentiation defects in the follicular cells (Castor downregulation) when *Dronc* expression was eliminated in large mutant clones that included the presumptive follicular stem cells (compare Supplementary Figs.1f with 1g). However, these clones were rarely recovered (11,4% (4/35); total number of clones analysed n=35 in 85 germaria) and only appeared after applying an experimental regime of repetitive heat-shocks^23^ that was against our purpose to investigate the functional requirement of *Dronc* under moderate stress. More importantly, the clones showing phenotypes had often compromised the expression of *Dronc* in both the somatic cells and germline (Supplementary Fig.1g), thus preventing us to extract unambiguous conclusions. To circumvent these technical limitations, we took advantage of a conditional null allele of *Dronc* generated in the laboratory through genome engineering^21,24^. This allele contains a wild-type *Dronc* cDNA flanked by FRT recombination sites (Supplementary Fig. 1h). After Flippase-mediated recombination, the permanent excision from the genome of FRT-rescue cassette can efficiently convert wild type cells into mutant^21,24^. Importantly, this allele conversion process does not require extreme heat-shock treatment and it can reproducibly be induced with temporal and spatial precision in specific cell populations by combining the *flippase* recombinase with the Gal4/UAS gene expression system. Downstream of the FRT-rescue cassette, we placed a transcriptional activator QF^25^, which facilitates the expression of any gene of interest under the physiological regulation of *Dronc* (hereafter *Dronc*^FRT-Dronc-FRT-QF^, Supplementary Fig. 1h)^21^. Capitalising on these features and using the *109-30*-Gal4 driver, we reproducibly eliminated the expression of *Dronc* in the follicular stem cells and their progeny^10^ (Supplementary Fig. 2a-c); all of the inspected germaria (100%, n=14) showed GFP signal (QUAS-*GFP*) in the follicular stem cells and abut prefollicular escort cells (Supplementary Fig. 2d). More importantly, this genetic manipulation significantly reduced the total number of follicular cells, and the proportion of follicular cells expressing the follicular differentiation marker Castor (Figs. 2a-d and Supplementary Fig. 2c). Expectedly, similar results were obtained eliminating *Dronc* expression in both escort and follicular cells using the *ptc*-Gal4^26^ (Fig 2c, 2d and Supplementary Figs. 2e-h). To determine whether these phenotypes were due to proliferation defects, we analysed the cell cycle profile of follicular cells with the marker termed Fly-Fucci^27^. This analysis revealed an increased proportion of follicular cells in S-phase at expense of cells in G0 in *Dronc* mutant conditions (Figs. 2e-g). These results were confirmed by assessing the incorporation of the S-phase marker EdU (Fig. 2h). Since the accumulation of follicular cells in S-phase did not lead to an excess of follicular cells but a reduction (Fig. 2c) and did not alter other stages of the cell cycle (Fig. 2g), we rationalised that Dronc deficiency appeared to slow down the transition through the S-phase and consequently the proliferation rate. Discarding a relevant contribution of cell death to our phenotypes, we also noticed that the number of TUNEL-positive follicular cells both in wildtype and *Dronc*-mutant conditions was comparable (Supplementary Fig. 2i). Importantly, the overexpression of *Dronc* also failed to cause excess apoptosis in our experimental conditions (Supplementary Fig. 2i). Complementarily, we confirmed the specificity of the phenotypes generated by the QF allele replacing this element with a Suntag-HA-Cherry peptide (*Dronc*^FRT-Dronc-FRT-suntag-HA-cherry^)^21^ (Supplementary Figs. 1h). The excision of the FRT-rescue-cassette in follicular cells and subsequent expression of Suntag-HA-Cherry instead of Dronc caused a reduction in the number of follicular cells and a Castor downregulation comparable to that of the QF allele (compare Fig. 2c, 2d with 3a, 3b). Collectively, these results indicated that non-apoptotic activation of *Dronc* facilitates the proliferation and differentiation of follicular stem cells and their progeny under moderate stress conditions.

**Fig. 2.**
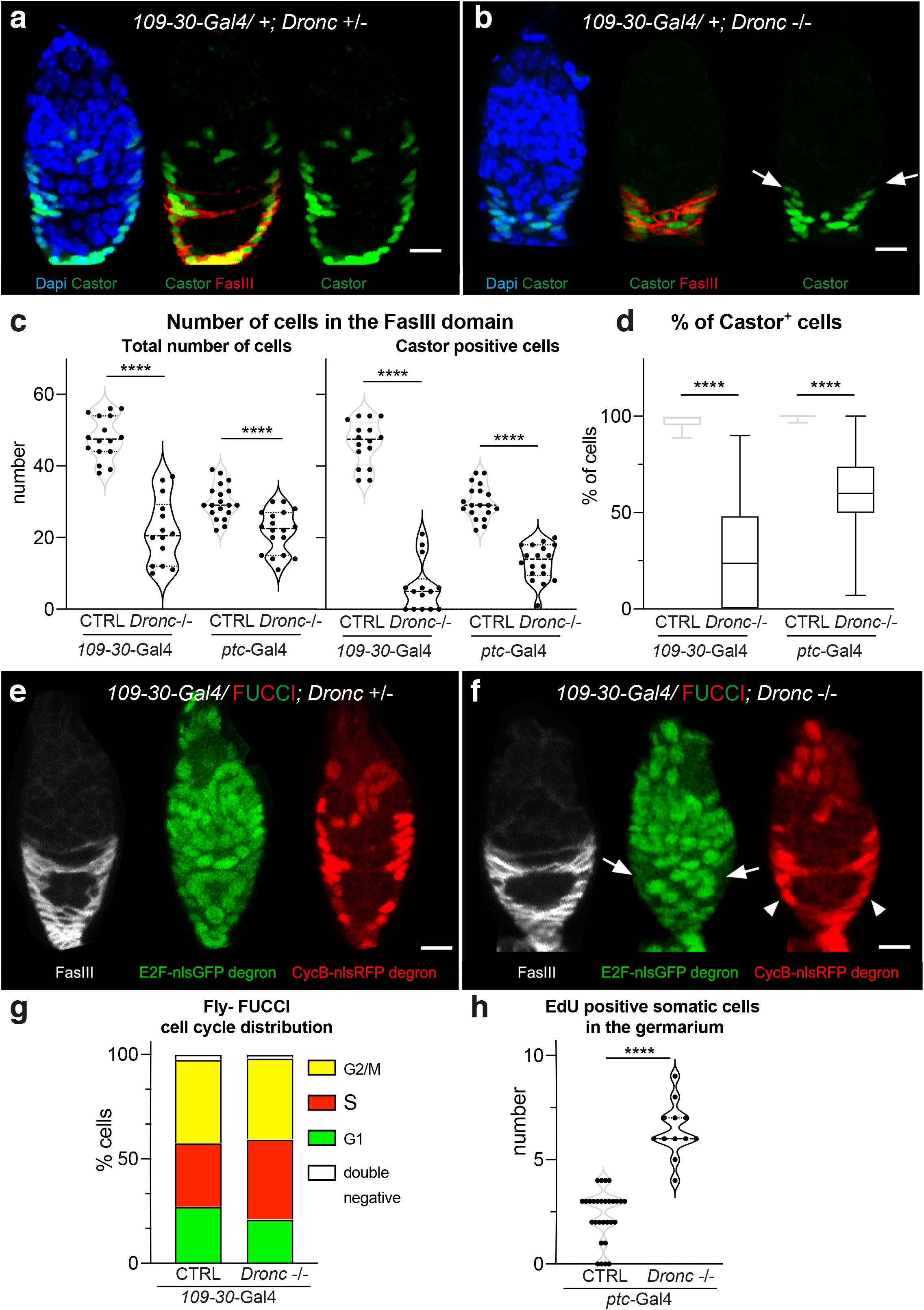
Functional characterization of *Dronc* in somatic cells. **a-b.** Confocal representative images comparing the expression of the follicular cell marker Castor in control (a: *109-30*-Gal4/+; *Dronc^KO^ Tub-G80^ts^*/+, n=16) versus mutant (b: *109-30*-Gal4/+; *Dronc^KO^ Tub-G80*^ts^/ UAS-*flippase Dronc^KO-FRT-Dronc-GFP-APEX-FRT-QF^*, n=14) germaria. Notice the reduction in the number of Castor-expressing cells in the follicular region (white arrows). Dapi labels DNA (blue); Castor (green); FasIII (red). Scale bars represents 10 μm. In the entire Figure, experimental flies were kept for 14 days at 29°C after eclosion and prior dissection. **c.** Quantification of total number of follicular cells (left) or Castor-expressing cells (right) within the FasIII cellular domain in either heterozygous or homozygous *Dronc* mutant cells generated using the *109-30*-Gal4 and *ptc*-Gal4 drivers, respectively; the *n* number for each column in order of appearance n=16, n=14, n=20, n=17, n=19, n=18. Statistical significance was established by using unpaired parametric T-test (****p≤0.001). Median and quartiles are shown in the violin plots of the entire Figure. d. Percentage of Castor-expressing cells versus the total number of Follicular cells in germaria of the genotypes indicated in c (FasIII^+^ cells). Data are expressed as box- and-whiskers plots, with min to max range as whiskers. Statistical significance was established by using Mann-Whitney test (****p≤0.0001). *n* number is shown in c. **e-f.** Representative confocal images showing Fly-FUCCI labelling in control (D: *109-30*-Gal4/FUCCI; *Dronc^KO^ Tub-G80^ts^*/+) and *Dronc* mutant (E: *109-30*-Gal4/FUCCI; *Dronc^KO^ Tub-G80^ts^* / UAS-*flippase Dronc^KO-FRT-Dronc-GFP-APEX-FRT-QF^*) follicular cells. FasIII staining (gray) is used as a reference to locate the follicular cells in the germarium; green signal labels G2, M and G1; red signal labels S, G2 and M. Notice the accumulation of cells in S-phase (red signal (arrowheads) without green (arrows) in the *Dronc* mutant condition). **g**. Graph showing the relative percentage of cells in different phases of the cell cycle with germaria of the genotypes described in D and E; control (left: CTRL: n=15) versus *Dron*c-/-(right: n=16) germaria. **h**. Quantification of somatic cells in S phases labelled by EdU incorporation in control (CTRL: *ptc*-Gal4/+; *Tub-G80^ts^*/+; n=30) versus mutant (*Dron*c-/-: *ptc*-Gal4/+; *Dronc^KO^Tub-G80^ts^* / UAS-*flippase Dronc^KO-FRT-Dronc-GFP-APEX-FRT-QF^*; n=12) germaria.

**Fig. 3.**
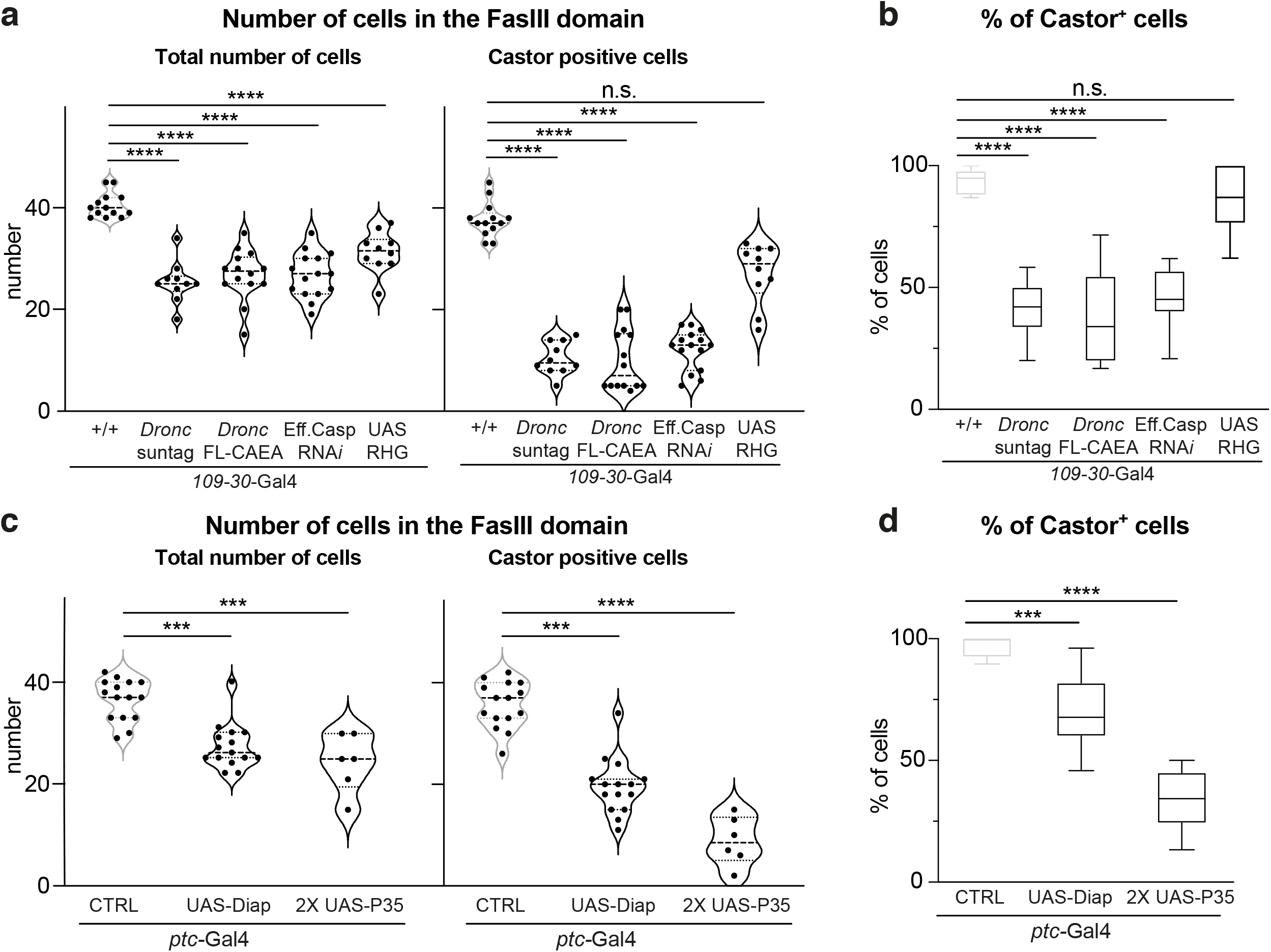
Analysis of molecular features required for Dronc to modulate stem cell follicular properties. **a.** Quantification of total number of follicular cells (left) or Castor-expressing cells (right) within the FasIII cellular domain after manipulating the expression of several members of the caspase pathway. The genotypes and the n number of the experiments are from left to right as follows: *109-30*-Gal4/+ (n=13); *109-30*-Gal4/+; *Dronc^KO^ Tub-G80^ts^* /UAS-*flippase Dronc*_*KO-FRT-Dronc-GFP-APEX-FRT*-suntag-Cherry-HA_ (n=15); *109-30*-Gal4/+; *Dronc^KO^ Tub-G80^ts^* / UAS-*flippase Dronc*^*KO-FRT-Dronc-GFP-APEX-FRT-Dronc*-FLCAEA^ (n=14); *109-30*-Gal4/UAS-*Drice*RNAi UAS-*Decay*RNAi; *Dronc*^KO^/UAS-*Dcp1*RNAi UAS-*Damm*RNAi (n=16); *109-30*-Gal4/UAS-*microRNA-RHG* (n=10); Statistical significance was established by using an ordinary one way ANOVA (****p≤0.0001) total number of cells and Kruskal-Wallis test, post test Dunn’s multiple comparison. **b.** Percentage of Castor-expressing cells versus the total number of Follicular cells in germaria of the genotypes indicated in a (FasIII^+^ cells). Data are expressed as box- and-whiskers plots, with min to max range as whiskers. Statistical significance was established by using ordinary one-way Anova test (****p≤0.0001; n.s.= p ≥ 0.5). *n* number is shown in a. **c.** Quantification of total number of follicular cells (left) or Castor-expressing cells (right) within the FasIII cellular domain in the following genotypes from left to right: *ptc*-Gal4/+ (n=15); *ptc*-Gal4/+; UAS-*Diap1* (n=15); *ptc*-Gal4/UAS-*P35; Dronc^KO^ Tub-G80^ts^* UAS-*P35* (n=6). Statistical significance was established by using Kruskal-Wallis test, post-test Dunn’s multiple comparison (****p≤0.0001; ***p≤0.001) **d.** Percentage of Castor-expressing cells versus the total number of Follicular cells in germaria of the genotypes indicated in c (FasIII^+^ cells). Data are expressed as box- and-whiskers plots, with min to max range as whiskers. Statistical significance was established by using Kruskal-Wallis test, post-test Dunn’s multiple comparison (****p≤0.0001; ***p≤0.001). *n* number is shown in c.

### The pro-survival effects of Dronc demands its catalytic activity

Most functions of caspases rely on their enzymatic activity, but some of the non-apoptotic roles only require protein-protein interactions^28,29^. To investigate the molecular activity of Dronc in follicular cells, we used a different conditional allele that contains after the FRT-rescue-cassette a mutant form of *Dronc* with two mutations, C318A and E352A (*Dronc*^FRT-Dronc-FRT-FLCAEA^; Supplementary Fig. 1h^21^). These mutations prevent the enzymatic function and proteolytic activation of *Dronc*, respectively^30,31^. The expression of Dronc^FLCAEA^ (*Dronc*^FRT-Dronc-FRT-FLCAEA^) in follicular cells reduced their number and caused Castor expression defects equivalent to that of other amorph *Dronc* alelles (Fig. 3a, 3b). Complementarily, the overexpression in somatic cells of the *Dronc* inhibitor *Diap-1 (Drosophila inhibitor of Apoptosis-1*)^32^ replicated the proliferation and differentiation phenotypes (Fig. 3c, 3d). These results strongly suggested an enzymatic requirement of *Dronc* in follicular cells. To further confirm this hypothesis, we examined the potential implication of the primary *Dronc* substrates in our cellular scenario; the so-called effector caspases (drICE, DCP-1, Decay and Damm)^33^. To avoid the functional redundancy between them^34^, we simultaneously expressed validated UAS-RNAi transgenes^33^ against all of them using the *109-30*-Gal4 driver (Fig. 3a, 3b). In parallel, we also conducted experiments overexpressing the effector caspase inhibitor P35^35^ (Fig. 3c, 3d). Both genetic manipulations mimicked the phenotypes obtained in *Dronc* mutant conditions (Figs. 3a-d). Finally, we assessed whether *Dronc* activation was linked to the pro-apoptotic factors Hid, Grim and Reaper by overexpressing a microRNA that simultaneously targets their expression. Interestingly, although this experiment also caused a reduction in the number of follicular cells (Fig. 3a, 3b), the proportion of Castor-expressing cells remained unaltered (Fig. 3b). This result led us to draw the important conclusion that the differentiation and proliferation defects caused by Dronc deficiency can be uncoupled. Furthermore, whereas Dronc requires the pro-apoptotic proteins to implement its pro-proliferative functions, these factors are largely dispensable for the differentiation process. Together, these experiments suggested that the non-apoptotic activation of the caspase pathway provides prosurvival cues that sustain the proliferation and differentiation of the follicular cells under moderate stress.

### Caspase activation promotes Hh-signalling acting upstream of *smoothened*

Hh-signalling deficiency due to the overexpression of *Ci*-RNAi in follicular stem cells causes phenotypes highly reminiscent to that of *Dronc* mutant conditions^6,36^ (compare Figs. 2a-d with Supplementary Figs. 3a-d). Comparably, the ectopic expression of Ptc^1130X^ in Hh-producing cells (terminal filament, escort cells, and prefollicular cells)^10^ under the regulation of *ptc*-Gal4^26^ prevented the proliferation and differentiation of follicular cells (Supplementary Figs. 3c, 3d); *ptc*^1130X^ encodes for a form of Ptc highly stable at the plasma membrane due to a low internalisation rate^37^, which efficiently and specifically prevents Hh-signalling in Hh-producing cells^38^. Taking into consideration the previous results, we sought to examine the activation levels of Hh-pathway in *Dronc*-mutant conditions. The insufficiency of *Dronc* in somatic cells reduced the expression levels of the active form of Cubitus interruptus (Ci-155^39^, Gli in mammals), as well as the transcription of the universal Hh-target gene, *ptc*^12^ (Figs. 4a-c). Importantly, similar Hh-signalling defects were detected in human ovarian cells with somatic origin (OVCAR-3) deficient in *caspase-9* (Figs. 4d, 4e). To functionally confirm the crosstalk between caspases and Hh-signalling, we attempted to rescue the *Dronc* mutant phenotypes by overexpressing either a constitutively active form of *smoothened* (*smo*) or Ci. The overexpression of any of these Hh-components in follicular cells mutant for *Dronc* restored the proliferation and Castor expression defects (Figs. 4f-i and Supplementary Figs. 3d, 3e). These data strongly suggested a crosstalk between caspases and Hh-pathway in ovarian somatic cells.

**Fig. 4.**
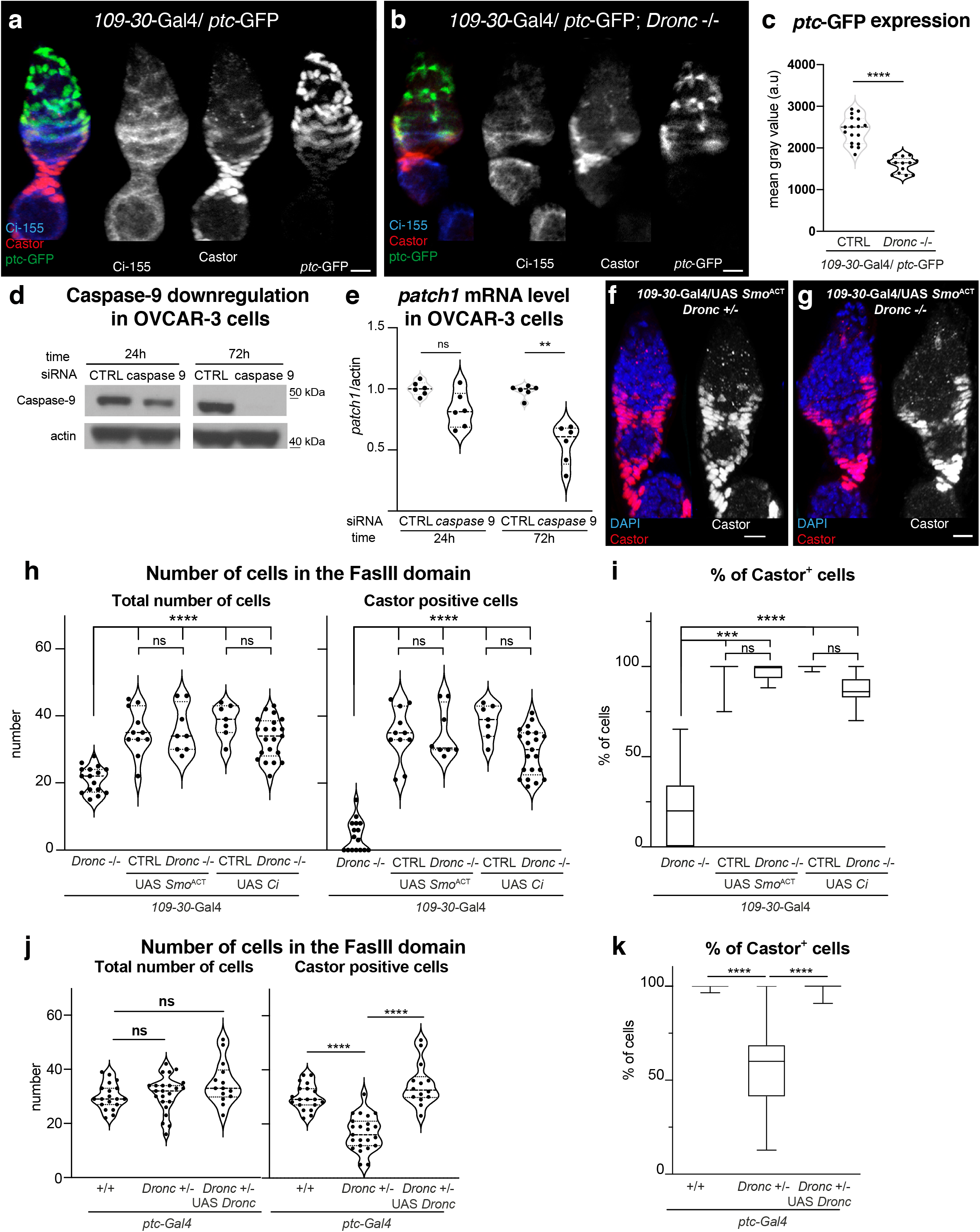
*Dronc* deficiency reduces Hh-signalling in *Drosophila* and OVCAR-3 ovarian somatic cells. **a-b.** Representative confocal images showing the expression of Ci-155 (blue and gray channels), *ptc*-GFP (*ptc*-GFP is a *bona-fide* transcriptional read out of Hh-pathway and weak hypomorph alelle^41^; green and gray channels) and Castor (red and gray channels) in either a control (A) or a *Dronc* mutant germaria (B). Experimental flies were kept for 14 days at 29°C after eclosion and prior dissection. **c.** Quantification of *ptc*-GFP expression in either a control (n=17) or a *Dronc* mutant (n=13) germaria; unpaired parametric T-Test was used to establish the statistical significance (**** p≤0.0001). Median and quartiles are shown in the violin plots of the entire Fig. Experimental flies were kept for 14 days at 29°C after eclosion and prior dissection. **d.** Western blot showing Caspase-9 expression (upper lane) and actin (bottom lane, loading control) in either control or *Caspase-9* deficient OVCAR-3 cells (24h and 72h post-transfection of an shRNA against *Caspase-9*). Notice the strong downregulation of Caspase-9 at 72h. **e.** mRNA levels of *patch1* measured by Q-PCR in either control or *Caspase-9* deficient OVCAR-3 cells; a Mann Whitney unpaired T-test was used to establish the statistical significance (** p≤0.01). **f-g.** Castor expression (red and gray channels) in either follicular cells heterozygous (F) or homozygous (G) for *Dronc* expressing a constitutively active form of *smo* under the regulation of *109-30*-Gal4 driver. Dapi staining labels the nuclei. Experimental flies were kept for 14 days at 29°C after eclosion and prior dissection. **h.** Quantification of total number of follicular cells (left) or Castor-expressing cells (right) in the following genotypes from left to right: *109-30*-Gal4/+; *Dronc^KO^ Tub-G80^ts^* /UAS-*flippase Dronc^KO-FRT-Dronc-GFP-APEX-FRT-QF^* (n=16); *109-30*-Gal4/UAS-*smo^Act^*; *Dronc^KO^ Tub-G80^ts^* /+ (n=11); *109-30*-Gal4/UAS-*smo^Act^*; *Dronc^KO^ Tub-G80^ts^*/UAS-*flippase Dronc^KO-FRT-Dronc-GFP-APEX-FRT-QF^* (n=8); *109-30*-Gal4/ UAS-*Ci*; *Dronc^KO^ Tub-G80^ts^*/+ (n=7); *109-30*-Gal4/UAS-*Ci*; *Dronc^KO^ Tub-G80^ts^*/UAS-*flippase Drond^KO-FRT-Dronc-GFP-APEX-FRT-QF^* (n=21). Scale bars represents 10 μm.). Statistical significance was established by using for the UAS-*smo^Act^* an ordinary one-way Anova (****p≤0.0001; n.s.= p ≥ 0.5) for total number of cells and a Kruskal-Wallis test, post-test Dunn’s multiple comparison (****p≤0.0001; ***p≤0.001; n.s.= p ≥ 0.5) for the rest of the data. Experimental and control flies were kept for 14 days at 29°C after eclosion and prior dissection. i. Percentage of Castor-expressing cells versus the total number of Follicular cells in germaria of the genotypes indicated in h (FasIII^+^ cells). Data are expressed as box- and-whiskers plots, with min to max range as whiskers. Statistical significance was established by using Kruskal-Wallis test, post-test Dunn’s multiple comparison (****p≤0.0001; ***p≤0.001). *n* number is shown in h. **j**. Quantification of total number of follicular cells (left) or Castor-expressing cells (right) within the FasIII cellular domain in the following genotypes from left to right: *ptc*-Gal4/+; Tub-*G80*^ts^ /+ (n=19); *ptc*-Gal4/+; *Dronc*^KO^ Tub-*G80*^ts^/+ (n=23); *ptc*-Gal4/+; *Dronc*^KO^ *Tub-G80^ts^*/UAS-*Dronc* (n=14). Scale bars represents 10 μm. Statistical significance was established by using an one-way ordinary ANOVA (n.s.= p ≥ 0.5; ****p≤0.0001). Experimental flies were kept for 7 days at 29°C after eclosion and prior dissection. **k.** Percentage of Castor-expressing cells versus the total number of Follicular cells in germaria of the genotypes indicated in j (FasIII^+^ cells). Data are expressed as box- and-whiskers plots, with min to max range as whiskers. Statistical significance was established by using Kruskal-Wallis test, post-test Dunn’s multiple comparison (****p≤0.0001; ***p≤0.001). *n* number is shown in j.

Since we rescued the mutant phenotypes of *Dronc* by either expressing an active form of Smo or Ci, we rationalised that the intersection of *Dronc* with the Hh-pathway might occur upstream of *smo*. To test this possibility, we performed classical genetic epistasis between *Dronc* and *ptc*. Interestingly, double heterozygous *ptc-Dronc* germaria (*ptc*-Gal4/+; *Dronc^KO^* /+; notice that the P-element insertion in the regulatory region that gave rise to *ptc*-Gal4 created a weak hypomorph allele^40^) showed a normal number of follicular cells; however, Castor expression was downregulated (Figs. 4j, 4k and Supplementary Fig. 4a). This result suggested a potential genetic interaction between *ptc* and *Dronc* and confirmed the possibility of uncoupling the differentiation and proliferation phenotypes (compare microRNA results in Fig. 3a, 3b with Figs. 4j, 4k). Furthermore, they showed that whereas the proliferation phenotypes require a full Dronc insufficiency (Figs, 2b, 2c), the Castor defects appear in double heterozygous conditions (Figs.3c, 3d). Importantly, the differentiation phenotypes in *ptc-Dronc* mutant flies were correlated with lower levels of Hh-signalling, as indicated by the downregulation of *ptc*-GFP (compare Fig. 4a, with 4b and Fig. Supplementary Fig. 4a; notice that *ptc*-GFP is also a *ptc* hypomorph allele^41^) and Ci-155 (compare Supplementary Fig. 4b). Confirming the legitimate nature of the potential genetic interaction between *ptc* and *Dronc*, Castor was also downregulated in ovarian somatic cells expressing a *Dronc*-RNAi construct, as well as in double heterozygous flies combining null alleles for *ptc*^S2^ and *Dronc^KO^* (*ptc*^S2^ /+; *Dronc*^KO^/+) (Supplementary Fig. 4c, 4d). Conversely, the overexpression of *Dronc* rescued the Castor expression defects in *ptc*-Gal4:*Dronc*^KO^ germaria (Figs. 4j, 4k). Together, these experiments confirmed a *bona fide* but initially counterintuitive genetic interaction between *ptc* and *Dronc*. Because Ptc is the receptor of Hh but acts as a negative regulator in terms of signalling, one would predict the phenotypic rescue of *Dronc* deficiency after reducing *ptc* expression. Instead, the double heterozygous *ptc-Dronc* germaria displayed phenotypes highly reminiscent to that of caused by either Hh-signalling or *Dronc* deprivation (compare Figs. 2c, 2d with 4j, 4k; compare Fig. 4b with Supplementary 4a).

### Dronc regulates Hh-signalling through the fine-tuning of Ptc

To better understand at the molecular level the interplay between *Dronc* and *ptc*, we investigated the Ptc protein levels in *ptc-Dronc* mutant germaria. Although *ptc* was transcriptionally downregulated (Figs. 4a-c and Supplementary Fig. 4a), the protein levels were strikingly elevated within escort and pre-follicular cells (Figs. 5a-c). Furthermore, the Ptc-positive punctae were significantly enlarged (Supplementary Fig. 4e). Interestingly, striking protein aggregates were also observed overexpressing Ptc^1130X^ in our mutant condition (Supplementary Figs. 4f-h). This result was particularly remarkable considering the low internalisation rate of Ptc^1130X 37^ (please, see discussion). To assess the biological significance of Ptc aggregates, we either overexpressed a *Ptc*-RNAi construct or used a combination of *ptc* alleles that further compromised its transcriptional activation. This experimental configuration was important to reduce the Ptc protein levels independently of the Ptc molecular activity. Both genetic manipulations largely restored the expression of Castor (Figs. 5d, 5e and Supplementary Fig. 4i). These results strongly suggested that *Dronc* takes part in the molecular network that regulates the fine-tuning of Ptc protein levels in ovarian somatic cells. Moreover, the accumulation of Ptc in caspase-deficient cells is largely responsible for the Hh-signalling deprivation and the phenotypes in follicular cells.

**Fig. 5.**
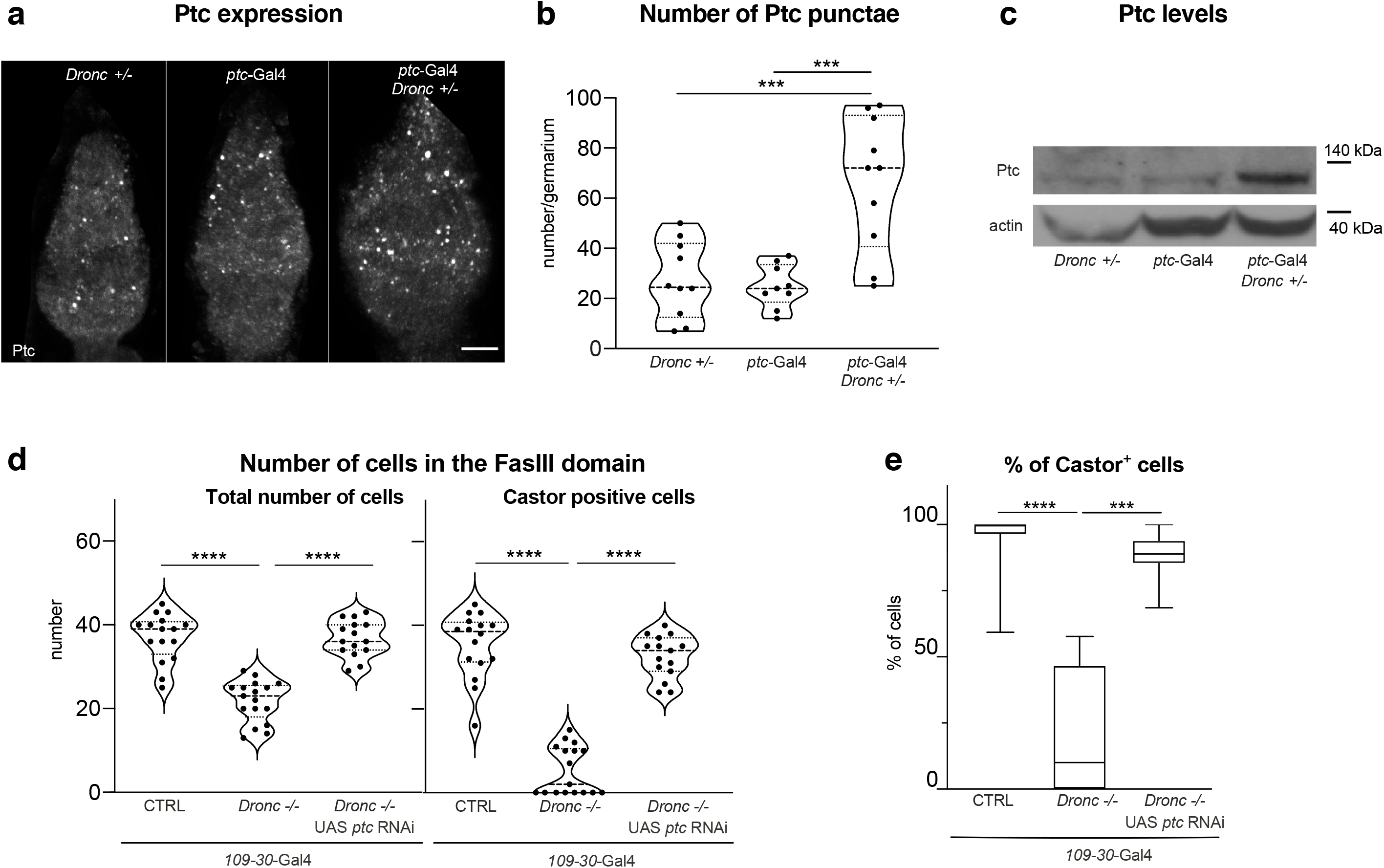
*Dronc* modulates Hh-signalling through the fine regulation of Ptc protein levels. **a.** Ptc immunostaining (gray channel) in germaria of the following genotypes: (*Dronc^KO^ Tub-G80^ts^* /+); (*ptc*-Gal4/+; *Tub-G80^ts^*/+); (*ptc*-Gal4/+; *Dronc^KO^ Tub-G80^ts^*/+). Scale bars represents 10 μm. In the entire Figure, experimental flies were kept for 7 days at 29°C after eclosion and prior dissection. **b.** Relative number of Ptc-positive punctae per germaria of the genotypes indicated in A: (*Dronc^KO^ Tub-G80^ts^* /+); (n=10); (*ptc*-Gal4/+; *Tub-G80^ts^*/+) (n=9); (*ptc*-Gal4/+; *Dronc^KO^ Tub-G80^ts^* /+). (n=10). A two-way ANOVA Tukey’s multiple comparisons test was used to establish the statistical significance (*** p≤0.001). Median and quartiles are shown in the violin plots of the entire Fig. **c.** Western blot showing Ptc (upper lane) and actin (bottom lane, loading control) expression in ovaries of the genotypes shown in A. Notice the Ptc accumulation in double heterozygous germaria (*ptc*-Gal4/+; *Dronc^KO^ Tub-G80^ts^* /+). **d.** Quantification of total number of follicular cells (left) or Castor-expressing cells (right) within the FasIII cellular domain in the following genotypes from left to right: *ptc*-Gal4/+; *Tub-G80^ts^*/+ (n=17); *ptc*-Gal4/+; *Dronc^KO^ Tub-G80^ts^* /UAS-*flippase Dronc^KO-FRT-Dronc-GFP-APEX-FRT-QF^* (n=16); *ptc*-Gal4/UAS-*ptc*-RNAi; *Dronc^KO^ Tub-G80^ts^* / UAS-*flippase Dronc^KO-FRT-Dronc-GFP-APEX-FRT-QF^* (n=15). Statistical significance was established by using an ordinary one-way ANOVA (**** p≤0.0001) for the total number of cells and a by using Kruskal-Wallis test, post-test Dunn’s multiple comparison (****p≤0.0001) for the castor positive cells. **e.** Percentage of Castor-expressing cells versus the total number of Follicular cells in germaria of the genotypes indicated in d (FasIII^+^ cells). Data are expressed as box- and-whiskers plots, with min to max range as whiskers. Statistical significance was established by using Kruskal-Wallis test, post-test Dunn’s multiple comparison (****p≤0.0001; ***p≤0.001) *n* number is shown in d.

### Dronc differentiation phenotypes are partially linked to Ptc-induced autophagy

In addition to the Hh-regulatory role, Ptc can induce autophagy in ovarian somatic cells from *Drosophila* and mammals^14,42^. Therefore, we investigated whether Ptc aggregates observed in our mutant conditions were able to induce autophagy. To assess the autophagy flux in the germarium, we used the expression of Ref2P (the *Drosophila* ortholog of the mammalian p62^43^). Whereas the activation of autophagy reduces the intracellular levels of Ref2P/p62, its inhibition facilitates Ref2P/p62 accumulation^44^. In agreement with the previous literature^14,42^, the expression of Ref2P increased in a genetic condition that modestly reduced Ptc protein levels (*ptc*-Gal4/+) (Figs. 6a-c). However, this upregulation was prevented halving the dose of *Dronc* (*ptc*-Gal4/+; *Droned*^KO^ /+) (Figs. 6a-c). These results suggested that Dronc deficiency can increase autophagy. To evaluate the contribution of autophagy to *Dronc* phenotypes, we compromised the expression of Atg1 using UAS-RNAi transgene with demonstrated activity in the ovary^45^. This genetic manipulation did not alter the number of follicular cells but largely rescued the Castor expression defects of *ptc-Dronc* germaria (Figs. 6d, 6e). These findings supported the hypothesis that non-apoptotic activation of *Dronc* modulates the intracellular levels of Ptc, which subsequently determine the fine-tuning of Hh-pathway and autophagy. Furthermore, they showed that the differentiation phenotypes induced by *Dronc* deficiency are strongly linked to the *Ptc*-dependent activation of autophagy. Next, we investigated the potential conservation of the *Drosophila* autophagy-related findings in human OVCAR-3 cells. Although the protein levels of p62 remained unaltered in *Caspase-9* mutant cells in standard culture conditions (Figs. 6f, 6g), they were significantly reduced after adding low concentrations of EtOH (Figs. 6f, 6g). Importantly, previous reports have shown that low levels of EtOH can trigger moderate cellular stress and activation of autophagy^46^. Confirming the specificity of p62 downregulation in our experiments, the inhibition of autophagy adding bafilomycin^47^ restored the expression levels of p62 in *Caspase-9* deficient cells treated with EtOH (Figs. 6f, 6g). These findings preliminarily suggest that the regulatory role of caspases on Hh-signalling and autophagy under moderate stress could be relevant in human cellular settings.

**Fig. 6.**
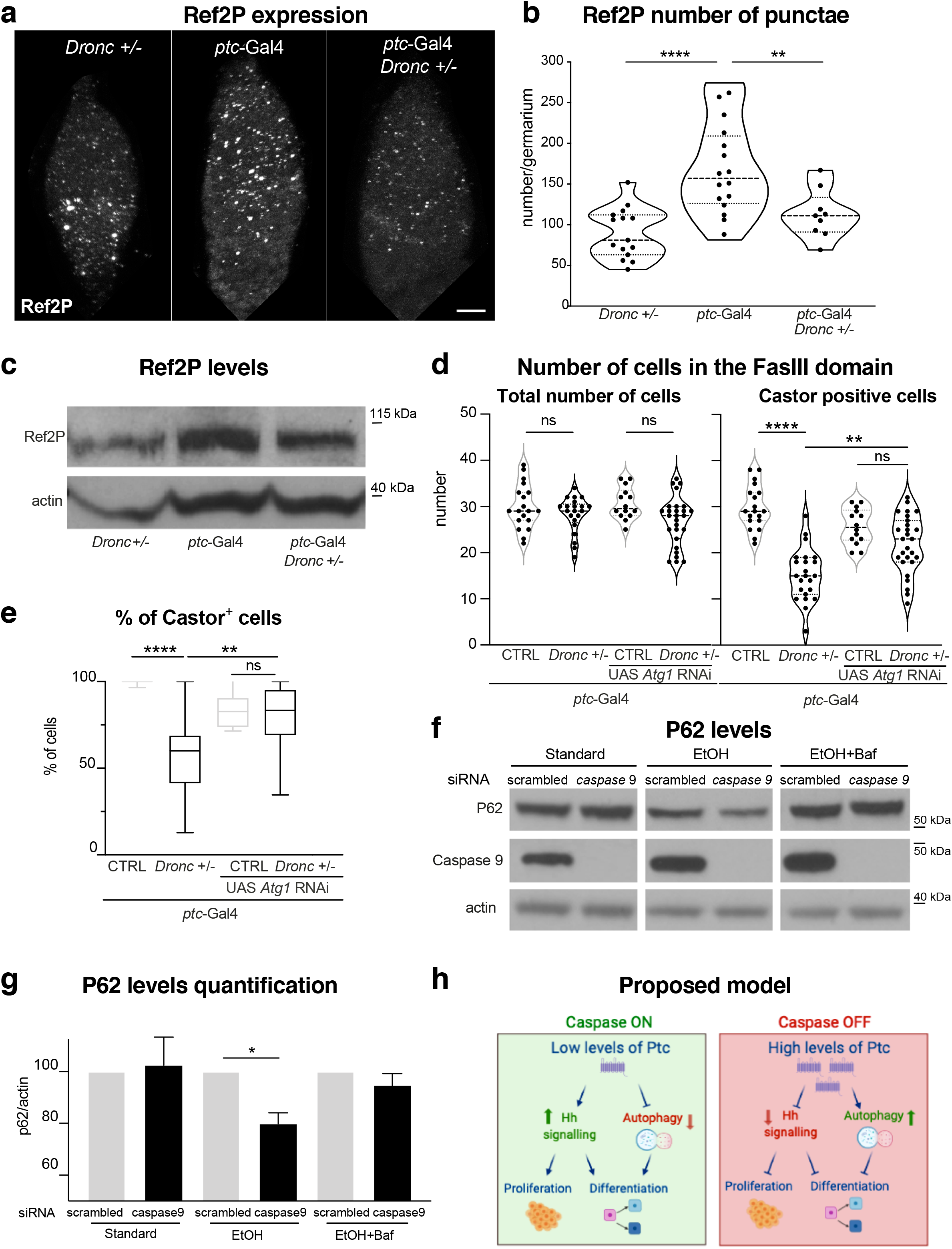
*Dronc* differentiation phenotypes are partially linked to Ptc-induced autophagy. **a.** Ref2P immunostaining (gray channel) in germaria of the following genotypes: (*Dronc^KO^ Tub-G80^ts^* /+); (*ptc*-Gal4/+; *Tub-G80^ts^*/+); (*ptc*-Gal4/+; *Dronc^KO^ Tub-G80^ts^*/+). Scale bars represents 10 μm. In the entire Figure, experimental flies were kept at 29°C after adult eclosion and prior dissection for 7 days **b.** Relative number of Ref2P-positive punctae per germaria of the genotypes indicated in A: (*Dronc^KO^ Tub-G80^ts^* /+); (n=15); (*ptc*-Gal4/+; *Tub-G80^ts^*/+); (n=16); (*ptc*-Gal4/+; *Dronc^KO^ Tub-G80^ts^* /+); (n=9). an ordinary one-way ANOVA Tukey’s multiple comparisons test was used to establish the statistical significance (** p≤0.01, ****p≤0.0001). **c**. Western blot showing Ref2P (upper lane) and actin (bottom lane, loading control) in ovaries of the genotypes shown in A. Notice the Ref2P reduction in double heterozygous germaria (*ptc*-Gal4/+; *Dronc^KO^ Tub-G80^ts^* /+) compared to the (*ptc*-Gal4/+; *Tub-G80^ts^*/+) control. **d.** Quantification of total number of follicular cells (left) or Castor-expressing cells (right) within the FasIII cellular domain in the following genotypes from left to right: *ptc*-Gal4/+; *Tub-G80^ts^*/+ (n=19); *ptc*-Gal4/+; *Dronc^KO^ Tub-G80^ts^*/+ (n=23); *ptc*-Gal4/UAS-*Atg*-RNAi; *Tub-G80^ts^* / + (n=14); *ptc*-Gal4/UAS-*Atg*-RNAi; *Dronc^KO^ Tub-G80^ts^* / + (n=27). Statistical significance was established by using an ordinary one way ANOVA test for total number of cells and an ordinary one-way Anova test, Tukey’s multiple comparison post-test for the castor positive cells (**** p≤0.0001, *** p≤0.001, n.s.= p ≥ 0.5). **e.** Percentage of Castor-expressing cells versus the total number of Follicular cells in germaria of the genotypes indicated in d (FasIII^+^ cells). Data are expressed as box- and-whiskers plots, with min to max range as whiskers. Statistical significance was established by using Kruskal-Wallis test, post-test Dunn’s multiple comparison (****p≤0.0001; **p≤0.01) *n* number is shown in d. **f.** Western blot showing the expression levels of the autophagy marker p62 (upper lane), Caspase-9 (middle lane) and Actin (bottom lane, loading control) in either scrambled or *Caspase-9* deficient OVCAR-3 cells; the protein levels of the different read outs were measured at 72h after siRNA treatment in cells grown during the last 4 h before sample processing in our standard cell culture conditions, in cell culture media containing EtOH (0.2%), and in cell culture media containing EtOH (0.2%) + bafilomycin A1 (400nM). **g.** Quantification of p62 protein levels in the experimental conditions described in E. one sample T Wilcoxon test was used to calculate statistical significance, * p≤0.05, n≥3. Bars indicate value of the mean while error bars represent the *Standard Deviation SD*. **h**. Model summarising the non-apoptotic caspase effects in ovarian somatic cells. Green and red colours indicate activation or silencing, respectively.

## DISCUSSION

Although caspases have traditionally been studied as main drivers of apoptosis, recent findings indicate the implication of these enzymes in the regulation of basic cellular functions independent of apoptosis. However, a complete understanding of such functions remains elusive in most cellular settings, including stem cells. Our findings strongly suggest that the non-apoptotic activation of the caspase-pathway is key to sustain Hh-signalling and prevent autophagy in ovarian somatic cells under moderate stress conditions. Furthermore, these unexpected caspases functions are crucial to facilitate the proliferation and differentiation of ovarian somatic cells in suboptimal cellular conditions. These findings shed light on unknown aspects of caspase biology, whilst conferring a pro-survival role to these enzymes.

### Non-apoptotic activation of Dronc acts as a pro-survival factor in ovarian somatic stem cells

Our experiments have shown widespread expression and activation of caspases in *Drosophila* ovarian somatic cells independent of apoptosis (Fig. 1 and Supplementary Fig. 1). Confirming the non-apoptotic nature of such caspase activation, caspase deficiency compromises the cell proliferation and differentiation of follicular stem cells and their progeny (Fig. 2) through the modulation of Hh-signalling (Fig. 4) and autophagy (Fig. 6). Furthermore, the differentiation phenotypes linked to *Dronc* mutant conditions appear to be largely independent of the expression of the pro-apoptotic factors and can occur without altering the number of follicular cells (microRNA-*RGH* data Fig. 3a, 3b). Together, these findings caution against the generic association of non-apoptotic patterns of caspase activation with the phenomenon of anastasis (pure recovery of caspase-activating cells from the “brink of death”)^48,49^. Conversely, they suggest that non-apoptotic caspase activation is essential for regulating cell signalling and pro-survival functions beyond apoptosis^1–3,50^

### Non-apoptotic caspase functions can involve the entire apoptotic pathway

Despite we have provided solid evidence indicating that non-apoptotic caspase activity is required to modulate the features of follicular cells under stress conditions, these functions still rely on the enzymatic activity of initiator and effector caspases (Fig. 3a). Furthermore, the caspase-dependent regulation of follicular cell proliferation also requires the expression of the conventional pro-apoptotic factors Hid, Grim and Reaper (Fig. 3a). These results indicate that not only caspases but the entire apoptotic pathway can be activated to implement non-apoptotic functions. Importantly, this is not exclusive of our *Drosophila* cellular model since a similar requirement has recently been demonstrated during the differentiation of mammalian muscle precursors^51^.

### Molecular basis of the caspase-dependent regulation of Hh-signalling and autophagy

At the molecular level, we provide evidence that non-apoptotic activation of Dronc prevents the accumulation of Ptc receptor (Fig. 5). Since Ptc accumulation is not correlated with its transcriptional upregulation (Fig. 4), we conclude that caspase activation likely enhances the degradation of Ptc. Supporting this hypothesis, a mutant form of Ptc^1130X^ that is rarely observed intracellularly due to its low rate of internalisation^37^ is strongly accumulated in large punctae in *Dronc*-mutant germaria (Supplementary Figs. S4f-h). However, two factors argue against the possibility that the enzymatic activity of *Dronc* directly facilitates the degradation of Ptc. First, the relevant targets implementing *Dronc* functions in our cellular model are the so-called effector caspases (Fig. 3). Second, the interplay between *Dronc* and *ptc* is highly specific to the *Drosophila* germarium, since caspases and Ptc coexist in many other *Drosophila* tissues without signs of interaction. An alternative explanation could be a potential regulatory action of caspases on the ubiquitin ligases involved in Ptc degradation. Interestingly, specific ubiquitin ligases have been shown to physically interact and activate Caspase-9 in mammalian cells deprived of Hh-signalling^52,53^. However, the direct connection of our caspase functions with Smurf (the ubiquitin ligase ortholog in *Drosophila*) also seems unlikely. If caspases would mediate the proteolytic degradation of Smurf, the excess of this protein in caspase mutant cells should reduce the Ptc levels^54^; instead, Ptc is significantly accumulated in caspase mutant conditions. Although further experiments out of the scope of this manuscript are needed to fully understand the molecular details of the relation between caspases and Ptc, our findings establish a novel functional connection between these two molecular factors, which in turn modulates essential cellular functions.

Beyond repressing Hh-signalling, the accumulation of Ptc in *Dronc* mutant follicular cells can induce autophagy^14,42^ (Fig. 6). Furthermore, the excess of autophagy contributes to the differentiation defects observed in *Dronc* mutant conditions (Fig. 6d, 6e). Previous studies have associated *Dronc* with the regulation of autophagy^55,56^; however, our data establish unprecedented links between this cellular process, Hh-pathway, and the caspases. Interestingly, as in *Drosophila* cells, *caspase-9* deficiency alter Hh-signalling (Fig. 4) and autophagy (Fig. 6) in human ovarian cells under moderate stress. These results preliminarily suggest that caspases could be part of an evolutionarily conserved genetic network able to modulate Hh-signalling and autophagy in ovarian somatic cells (Fig. 6h).

### Cellular, physiological and evolutionary implications of non-apoptotic caspase activation in ovarian somatic cells

At the cellular level, the proliferation phenotypes caused by caspase deficiency are likely correlated with Hh-signalling since solid evidence indicates the implication of this pathway in the regulation of the cell cycle^57,58^. Supporting this hypothesis, we have shown that the proliferation phenotypes disappear after restoring Hh-signalling in caspase mutant follicular cells (Fig. 4h, 4i). Independently, we have also observed that caspase-dependent proliferation and differentiation phenotypes can be uncoupled and require different Dronc protein levels. Whereas the downregulation of Castor appears in *ptc-Dronc* heterozygous cells (Fig. 4j), the proliferation defects only emerge in *Dronc* homozygous conditions (Fig. 2c). Interestingly, the expression of Castor and therefore the differentiation defects can be restored in our mutant conditions not only by Hh-signaling (Fig. 4h) but also by preventing the excess autophagy (Fig. 6d). These findings suggest that the differentiation process of follicular cells is coordinated by the opposing activities of Hh-signalling and autophagy. Additionally, they indicate that the Castor downregulation is not directly responsible for the proliferation defects (Fig. 6h). Taking together, our results support the hypothesis that non-apoptotic levels of caspase activation could be at the forefront of the cell survival mechanisms against cellular stress (Fig. 6h) but sustained caspase activation due to persistent signalling defects and/or environmental stress lead to apoptosis. This dual role of caspases coupled to different signalling pathways and cellular contexts appears to ensure proper development and cells homeostasis^59,60^ (our data), as well as the execution of cellular selection mechanisms^61^.

From a physiological perspective, it is has been reported that Hh-downregulation triggered by environmental stress restricts egg laying and promotes autophagy in *Drosophila*^62–64^. Similarly, Hh deregulation and/or exacerbated autophagy can compromise follicular development in mammalian systems^65,66^. Our work suggests that sublethal caspase activation influences Hh-signalling and autophagy (Fig. 6h), and therefore it might be part of a complex adaptive system that ensures timely egg maturation in stress situations.

Taking into consideration the non-apoptotic roles of ancient members of the caspase family^3,67,68^, our findings may also have evolutionary implications. Since Dronc can play a pro-survival role in somatic cells, our data support the hypothesis that caspases could initially sustain basic cellular processes, and only their inadvertent/persistent activation would lead to cell death^67^. From this perspective, these pro-apoptotic enzymes could act as pro-survival factors, thus inverting the widely held view regarding their most primitive function.

## MATERIAL AND METHODS

### Fly Strains and fly husbandry details

All fly strains used are described at www.flybase.bio.indiana.edu unless otherwise indicated. After 24h of egg laying at 25°C, experimental specimens were raised at 18°C, thus enabling the repression of Gal4 activity through a Gal80^ts^. This prevents lethality in our experiments during larval and pupal stages. After hatching, adults were then transferred from 18°C to 29°C until dissection time. At 29°C the repression of Gal80^ts^ disappears, and therefore gene expression via Gal4 is elicited within specific cell subpopulations of the germarium. The temperature shift of adult flies at 29°C was also maintained for those genetic combinations that were not lethal in previous developmental stages.

### Detailed Genotype Description

Full description of experimental genotypes appearing in each Figure.

**Fig. 1**

**1b**. Actin *DBS-S-QF*, UAS-*mCD8-GFP*, QUAS-*tomato-HA*/+;; QUAS-*Gal4*/+

**1c** and **1d**. Actin *DBS-S-QF*, UAS-*mCD8-GFP*, QUAS-*tomato-HA*/+; QUAS-*flippase* (BL30126)/+; Actin5C FRT-*stop*-FRT *lacZ*-nls/+ (BL6355)

**1f**. w;; *Dronc*^KO-Gal4^ / UAS-*Histone-RFP* (BL56555)

**1g.** w;; *Dronc*^TurboID^ (a gift from Masayuki Miura)/ Tm3, Sb

**Fig. 2**

**2a**: *109-30-Gal4* (BL7023)/+; *Dronc*^KO^ Tub-*G80^ts^* (BL7019)/+;

**2b**: *109-30-Gal4* (BL7023)/+; *Dronc*^KO^ Tub-*G80*^ts^ (BL7019)/ UAS-*flippase* (BL8209)

*Dronc*^KO-FRT-Dronc-GFP-APEX-FRT-QF^.

**2c, 2d.** From left to right:

*CTRL= 109-30-Gal4* (BL7023)/+; *Dronc*^KO^ Tub-*G80^ts^* (BL7019)/+;

*Dronc -/- = 109-30-Gal4* (BL7023)/+; *Dronc*^KO^ Tub-*G80*^ts^ (BL7019)/ UAS-*flippase* (BL8209) *Dronc*^KO-FRT-Dronc-GFP-APEX-FRT-QF^.

*CTRL=ptc-Gal4* (BL2017)/+; +/+

*Dronc -/- = ptc-Gal4* (BL2017)/+; *Dronc*^KO^ Tub-*G80*^ts^ (BL7019)/ UAS-*flippase* (BL8209) *Dronc*^KO-FRT-Dronc-GFP-APEX-FRT-QF^.

**2e.** *109-30-Gal4* (BL7023)/FUCCI(BL55123); *Dronc*^KO^ Tub-*G80*^ts^ (BL7019)/ +

**2f.** *109-30-Gal4* (BL7023)/FUCCI(BL55123); *Dronc*^KO^ Tub-*G80*^ts^ (BL7019) / UAS-*flippase* (BL8209) *Dronc*^KO-FRT-Dronc-GFP-APEX-FRT-QF^.

**2g.** From left to right:

*CTRL= 109-30-Gal4* (BL7023)/FUCCI(BL55123); *Dronc*^KO^ Tub-*G80*^ts^ (BL7019)/ +

*Dronc -/- = 109-30-Gal4* (BL7023)/FUCCI(BL55123); *Dronc*^KO^ Tub-*G80*^ts^ (BL7019) / UAS-*flippase* (BL8209) *Dronc*^KO-FRT-Dronc-GFP-APEX-FRT-QF^.

**2h.** From left to right:

*CTRL= ptc-Gal4* (BL2017)/+; Tub-*G80*^ts^ (BL7019)/+

*Dronc -/- = ptc-Gal4* (BL2017)/+; *Dronc*^KO^ Tub-*G80*^ts^ (BL7019) / UAS-*flippase* (BL8209) *Dronc*^KO-FRT-Dronc-GFP-APEX-FRT-QF^.

**Fig. 3**

**3a, 3b.** From left to right:

*+/+ = 109-30-Gal4* (BL7023)/+; *+* / +

*Dronc Suntag = 109-30-Gal4* (BL7023)/+; *Dronc*^KO^ Tub-*G80*^ts^ (BL7019) / UAS-*flippase* (BL8209) *Dronc*^*KO*-FRT Dronc-GFP-Apex FRT-Suntag-HA-Cherry^

*Dronc FLCAEA= 109-30-Gal4* (BL7023)/+; *Dronc*^KO^ Tub-*G80*^ts^ (BL7019) / UAS-*flippase* (BL8209) *Dronc*^*KO*-FRT Dronc-GFP-Apex FRT-Dronc FL-CAEA-Suntag-HA-Cherry^

*Eff. Casp. RNAi = 109-30-Gal4* (BL7023)/UAS-*Drice*RNAi UAS-*Decay*RNAi (a gift from Pascal Meier); UAS-*Damm*RNAi, UAS-*Dcp1*RNAi (a gift from Pascal Meier).

*UAS-RGH = 109-30-Gal4* (BL7023)/UAS-microRNA-RHG (a gift from Istwar Hariharan)

**3c, 3d.** From left to right:

*CTRL= ptc-Gal4* (BL2017)/

*UAS-Diap1 = ptc-Gal4* (BL2017)/UAS-*Diap1* (BL63819)

*2x P35 = ptc-Gal4* (BL2017)/UAS-P35 (BL5072); UAS-P35 (BL5073)/+

**Fig. 4**

**4a.** *109-30-Gal4* (BL7023)/*ptc*-GFP^CB02030^ (a gift from Isabel Guerrero). *ptc*-GFP^CB02030^ contains a P-element insertion in the promoter region of *ptc* that generates a weak hypomorph allele of *ptc*^41^.

**4b**. *109-30-Gal4* (BL7023)/*ptc*-GFP^CB02030^; *Dronc*^KO^Tub-*G80*^ts^ (BL7019)/ UAS-*flippase Dronc^KO-FRT-Dronc-GFP-APEX-FRT-QF^*

**4f.** *109-30-Gal4* (BL7023)/UAS-*smo*^Act^ (BL44621); *Dronc*^KO^Tub-*G80*^ts^ (BL7019)/+

**4g.** *109-30-Gal4* (BL7023)/UAS-*smo*^Act^ (BL44621); *Dronc*^KO^Tub-*G80*^ts^ (BL7019)/ UAS-*flippase* (BL8209) *Dronc^KO-FRT-Dronc-GFP-APEX-FRT-QF^*

**4h, 4i.** From left to right:

*Dronc -/- = CTRL= 109-30-Gal4* (BL7023)/; *Dronc*^KO^Tub-*G80*^ts^ (BL7019)/ UAS-*flippase* (BL8209) *Dronc^KO-FRT-Dronc-GFP-APEX-FRT-QF^*

*CTRL= 109-30-Gal4* (BL7023)/UAS-*smo*^Act^ (BL44621); *Dronc*^KO^Tub-*G80*^ts^ (BL7019)/+

*Dronc -/- = 109-30-Gal4* (BL7023)/UAS-*smo*^Act^ (BL44621); *Dronc*^KO^Tub-*G80*^ts^ (BL7019)/ UAS-*flippase* (BL8209) *Dronc^KO-FRT-Dronc-GFP-APEX-FRT-QF^*

*CTRL= 109-30-Gal4* (BL7023)/UAS-*Ci* (BL28984); *Dronc*^KO^Tub-*G80*^ts^ (BL7019)/+

*Dronc -/- = 109-30-Gal4* (BL7023)/UAS-UAS-*Ci* (BL28984); *Dronc*^KO^Tub-*G80*^ts^(BL7019)/ UAS-*flippase* (BL8209) *Dronc^KO-FRT-Dronc-GFP-APEX-FRT-QF^*

**4j, 4k.** From left to right:

*Dronc +/+ = ptc-Gal4* (BL2017)/+; Tub-*G80*^ts^ (BL7019)

*Dronc +/- = ptc*-Gal4 (BL2017)/+; *Dronc*^KO^Tub-*G80*^ts^ (BL7019)/+

*Dronc +/- UAS-Dronc = ptc*-Gal4 (BL2017)/+; *Dronc*^KO^Tub-*G80*^ts^ (BL7019)/UAS-*Dronc* (BL56198)

**Fig. 5**

**5a-c.** From left to right:

*Dronc*^KO^Tub-*G80*^ts^ (BL7019)/+

*ptc*-Gal4 (BL2017)/+; Tub-*G80*^ts^ (BL7019)/+

*ptc*-Gal4 (BL2017)/+; *Dronc*^KO^Tub-*G80*^ts^ (BL7019)/+

**5d.** From left to right:

*CTRL = 109-30-Gal4* (BL7023)/+; *Dronc*^KO^Tub-*G80*^ts^ (BL7019)/+

*Dronc* -/- = *109-30Gal4* (BL7023)/+; *Dronc*^KO^Tub-*G80*^ts^ (BL7019)/ UAS-*flippase* (BL8209) *Dronc^KO-FRT-Dronc-GFP-APEX-FRT-QF^*

*Dronc* -/- UAS-*ptc*-RNAi = *109-30Gal4* (BL7023)/UAS-*ptc*-RNAi (BL55686);

*Dronc*^KO^Tub-*G80*^ts^ (BL7019)/ UAS-*flippase* (BL8209) *Dronc^KO-FRT-Dronc-GFP-APEX-FRT-QF^*

**Fig. 6**

**6a-c.** From left to right:

*Dronc*^KO^Tub-*G80*^ts^ (BL7019)/+

*ptc*-Gal4 (BL2017)/+; Tub-*G80*^ts^ (BL7019)/+

*ptc*-Gal4 (BL2017)/+; *Dronc*^KO^Tub-*G80*^ts^ (BL7019)/+

**6d, 6e.** From left to right:

CTRL= *ptc*-Gal4 (BL2017)/+; Tub-*G80*^ts^ (BL7019)/+

*Dronc* +/- = *ptc*-Gal4 (BL2017)/+; *Dronc*^KO^Tub-*G80*^ts^ (BL7019)/+

CTRL= *ptc*-Gal4 (BL2017)/+; Tub-*G80*^ts^ (BL7019)/UAS-*Atg1*-RNAi (BL35177)

*Dronc* +/- = *ptc*-Gal4 (BL2017)/+; *Dronc*^KO^Tub-*G80*^ts^ (BL7019)/UAS-*Atg1*-RNAi (BL35177)

### Immunohistochemistry

Adult *Drosophila* ovaries were dissected on ice-cold PBS. Immunostainings and washes were performed according to standard protocols (fixing in PBS 4% paraformaldehyde, washing in PBT 0.3% (0.3% Triton X-100 in PBS). Primary antibodies used in our experiments were: anti-Castor (1:2000; a gift from Alex Gould); rabbit anti-HA (1:1000; Cell Signaling C29F4); mouse anti-β-Gal (1:500; Promega Z378B); chicken Anti-βGal (1:200, Abcam AB9361); Anti-FasIII (1:75, Hybridoma Bank 7G10); Anti-Ci-155-full length (1:50, Hybridoma Bank 2A1); Anti-Ptc (1:50, Hybridoma Bank Apa1); Anti-Ref2P (1:300, abcam 178440). Conjugated secondary antibodies (Molecular Probes) were diluted in 0.3% PBT and used in a final concentration (1:200): conjugated donkey anti-rabbit Alexa-Fluor-488 (A21206) or 555 (A31572) or 647 (A31573), conjugated donkey anti-mouse Alexa-Fluor-488 (A21202) or 555 (A31570) or 647 (A31571), conjugated goat anti-rat Life Technologies (Paisley, UK) Alexa-Fluor-488 (A21247) or 555 (A21434). The detection of biotinylated proteins was made using Streptavidin conjugated with the 488 fluorophore (1:500; S11223). Dapi was added to the solution with the secondary antibodies for labelling the nuclei (1:1000; Thermo Scientific 62248). Following incubation in secondary antibodies, samples were washed several times during 60 minutes in PBT. Finally, they were mounted on Poly-Prep Slides (P0425-72EA, Sigma) in Aqua-Poly/Mount (Polysciences, Inc (18606)).

### TUNEL staining

Like in the immunochemistry, follicles from adult *Drosophila* females were dissected in ice-cold PBS and fixed in PBS containing 4% formaldehyde for 20’. After fixation, the samples were washed 3 times for 15’ with PBS and subsequently permeabilised with PBS containing 0,3% triton and 0,1% sodium citrate for 8’ on ice. 3 PBS washes for 20’ with were performed also after permeabilisation. The *in situ* detection of fragmented genomic DNA was performed according to the DeadEnd colorimetric TUNEL (Terminal transferase-mediated dUTP nick-end labeling) system (Promega). Briefly, samples were first equilibrated at room temperature in equilibration buffer (5-10’) and then incubated with TdT reaction mix for 1 hour at 37°C in a humidified chamber to obtain the 3’-end labelling of fragmented DNA. The reaction was terminated with 3 washes for 15’ in PBS. If necessary, the TUNEL protocol was followed by standard immunofluorescent staining. The detection of TUNEL-positive cells was achieved by an incubation of 45’ with streptavidin-fluorophore conjugated dyes.

### EdU Staining

Adult female ovaries were dissected in PBS1X, transferred to a microfuge tube containing 10mM EdU in PBS1X and kept at room temperature on a shaker for 1□h. Ovarioles were then dissociated, fixed, and stained with primary and secondary antibodies as described above. The EdU detection reaction was performed according to the manufacturer’s manual (Thermo Fisher Scientific, C10640).

### Morphogenetic mosaics generation

Two-day old adult females of the genotype yw hs-*Flp*^1.22^/+; UAS-*flippase*/+; FRT80, *Dronc*^l29^/FRT80 Ubiquitin-*GFP* were given either two or four 1 hour heat-shocks at 37°C spread over 2 days (12h apart). This allowed variable mitotic recombination efficiency and therefore different number of genetic mosaics. The higher is the number of heat-shocks, the larger is the probability of covering a large fraction of tissue with mutant cells. After the last heat shock, flies were kept at 29°C under a regime of frequent transfer (every two days) to a fresh vial with standard food supplemented with yeast. Flies were dissected and immunostained 7days after the last heat shock.

### Imaging

*Drosophila* ovarioles were imaged using the Olympus Fluoview FV1200 and associated software. Z-stacks were taken with a 40X objective at intervals along the apical-basal axis that ensured adequate resolution along Z-axis (step size 0.5-1.5-μm). The same confocal settings were used during the image acquisition process of experimental and control samples. Acquired images were processed using ImageJ 1.52n, Adobe Photoshop2020 and Adobe Illustrator2020 in order to complete the Fig. preparation. When confocal images were rotated, a dark rectangular background was added to create regularly shaped figures.

### Image quantification

All of the images used in this study were randomised and blindly scored during the quantification process. Images for quantification purposes were processed with ImageJ 1.52p.

The total number of cells expressing either FasIII or UAS-Histone-RFP in the germarium was manually quantified in each different focal plane of the germarium using an ImageJ Cell Counter macro specifically written to that purpose. This macro avoids the duplicated counting of the same object in the different focal planes of the image of interest. The same procedure was followed to estimate the number of Castor-expressing cells in the follicular region of the germarium. The percentage of Castor-expressing cells in the follicular region was estimated dividing the total number of Castor-expressing cells by the total number of FasIII/UASHistone-RFP positive cells in the follicular region.

To quantify the number and size of Ptc and Ref2P-positive particles in the regions 1, 2a and 2b of the germarium (Fig.s 4B, 5B, and Fig. S4E), we first made a maximum projection of the total focal planes. Then we sequentially applied the thresholding and “Analyse Particles” plug-ins from ImageJ. An equivalent image processing method was used to estimate the Ptc expression levels in Fig.s 3C and Fig. S4H. The “mean gray value” function of image J was used in this instance to estimate the GFP levels.

### Western Blot

Adult *Drosophila* ovaries were dissected in ice-cold PBS and snap-frozen in liquid nitrogen. Subsequently, they were homogenised in NP40 buffer [150 mM NaCl, 50 mM Tris-HCl pH 7.5, 5% glycerol, 1% IGEPAL CA-630]. Cells were harvested using trypsin/EDTA and centrifuged at 300g for 5’. Pellets were washed in PBS and then treated with RIPA lysis buffer 1x [150 mM NaCl, 50 mM Tris-HCl pH 7.5, 0.1 mM EGTA, 0,5 mM EDTA, 1% Triton X-100]. Halt Protease and Phosphatase Inhibitor Cocktail (Thermo Scientific Pierce) and Benzonase (BaseMuncher, Expedeon) were added according to the manufacturer’s instructions. Protein content was determined using Bradford reagent (Bio-Rad). Extracts were mixed with NuPAGE LDS Sample Buffer and separated by SDS-PAGE. For performing the SDS-PAGE electrophoresis, lysates were loaded and run in NuPAGE Bis-Tris Gels in NuPAGE MOPS SDS Running Buffer (Thermofisher Scientific). Protein blot transfers were performed using Trans-Blot Turbo Transfer System (Biorad). Nitrocellulose blots were incubated at room temperature for 30’ in blocking buffer [Tris-buffered saline with 0.1% Tween containing 5% non-fat dried milk] and then incubated overnight at 4°C in the same blocking solution with the corresponding antibodies. After washing three times for 15’ each with Tris-buffered saline containing 0.1% Tween, the blots were incubated with horseradish peroxidase-conjugated (HRP) IgG, followed by washing. Immunoreactive bands were detected using the SuperSignal West Pico PLUS Chemiluminescent Substrate (Thermofisher Scientific). Developed CL-XPosure films (Thermofisher Scientific) were scanned using a flat-bed scanner and the density of the bands was measured using Gel Analyzer plugin in ImageJ software. Primary antibodies used: Anti-Ptc (1:500, Hybridoma Bank Apa1); Anti-Ref2P (1:500, abcam 178440); Anti-Actin (1:500, Hybridoma Bank JLA20s); Anti-Ci-155-full length (1:500, Hybridoma Bank 2A1); Anti-Caspase-9 (C9) (1:1000, Cell Signalling 9508); Anti-ß-Actin-Peroxidase (1:20000, Sigma A3854), Anti SQSTM1 / P62 antibody (1:5000, GeneTex GTX111393).

### Cell culture mammalian cells

OVCAR-3 cells were maintained in RPMI (Sigma, R8758), supplemented with 10% FBS (Life Technologies, 10500064) and grown at 37°C in a humidified atmosphere with 5% CO_2_. For the experiment shown in Fig. 5c and 5d, we replaced the media with fresh media containing either EtOH (0.2%) or EtOH (0.2%) + the inhibitor of autophagy bafilomycin A1 (400nM, Merck Chemicals). Cells were grown in these two different cell culture media during the last 4 hours previous the sample processing.

### RNA interference

Small interfering RNA (siRNA) specific for Caspase-9 (ON-TARGETplus SMART pool human L-003309-00-0005, 842), PTCH1 (ON-TARGETplus Human PTCH1, L-003924-00-0005, 5727) and non-targeting controls (ON-TARGET plus Non-targeting Pool, D-001810-10-05) were purchased from Dharmacon Inc. (UK). Cells were plated and transfected the day after with Oligofectamine™ Transfection Reagent (Thermofisher 12252) in the presence of siRNAs according to the manufacturer’s instructions. Cells were kept in the transfection mix before processing for western blot or Q-PCR at the specified time points (24h and 72h).

### Gene expression analyses by Q-PCR

RNA extraction was performed using the Qiagen RNeasy Plus kit (74034). cDNAs were synthesised with Maxima First Strand cDNA synthesis kit (Molecular Biology, Thermofisher, K1642) Q-PCR were performed using QuantiNova SYBR Green PCR Kit (Qiagen, 208054). Detection was performed using Rotor-Gene Q Real-time PCR cycler (Qiagen).

Data was analysed using the Pfaffl method, based on ΔΔ-Ct and normalised to actin as the housekeeping gene.

Gene expression was estimated with the following primers:

*Patched1*:

Forward CCACGACAAAGCCGACTACAT
Reverse GCTGCAGATGGTCCTTACTTTTTC

*B-actin:*

Forward CCTGGCACCCAGCACAAT
Reverse GGGCCGGACTCGTCATAC.

## CONTRIBUTIONS

L.A.B-L. was responsible for the initial conception of the work and original writing of the manuscript. The experimental design was elaborated by A.G and L.A.B-L. A.G was responsible for most of the experimental work. D.I. performed some experiments under the supervision of A.G. The Fig. preparation was made by A.G and L.A.B-L. All co-authors have provided useful criticisms and commented on the manuscript before submission.

## ACKNOWLEDGEMENTS

Thanks for providing flies and reagents to; Isabel Guerrero (*ptc-GFP*; Centro de Biología Molecular); Pascal Meier (UAS-*Dronc*-RNAi, UAS-*Drice*-RNAi, UAS-*Dcp*-RNAi, UAS-*Damm*-RNAi and UAS-*Decay*-RNAi); Iswar Hariharan (UAS-*miRGH*); Alex Gould (anti-Castor antibody, CRICK Institute), Masayuki Miura (*Dronc*^*T*urboID^ and *Drlce*^TurboID^), the Developmental Studies Hybridoma Bank (antibodies), Addgene (*pCDNA3-connexin-GFP-Apex2* plasmid), Bloomington Stock Center (fly strains), Kyoto Stock Center (fly strains), and DGRC (wild-type cDNA of *dronc*). Thanks to Genewiz and Bestgene for making the DNA synthesis and generating transgenic flies, respectively. Thanks also to Ulrike Gruneberg, Sonia Muliyil, Xavier Franch-Marro, Jordan Raff and the caspaselab members (https://www.caspaselab.com) for the critical reading of the manuscript and valuable suggestions. This work has been supported by Cancer Research UK C49979/A17516 and the John Fell Fund from the University of Oxford 162/001. L.A.B-L. is a CRUK Career Development Fellow (C49979/A17516) and an Oriel College Hayward Fellow. A.G. is a postodoctoral fellow of CRUK (C49979/A17516).

## REFERENCES

1 Aram, L., Yacobi-Sharon, K. & Arama, E. CDPs: caspase-dependent non-lethal cellular processes. Cell Death Differ 24, 1307–1310, doi:10.1038/cdd.2017.111 (2017).

2 Baena-Lopez, L. A. All about the caspase-dependent functions without cell death. Semin Cell Dev Biol, doi:10.1016/j.semcdb.2018.01.005 (2018).

3 Bell, R. A. V. & Megeney, L. A. Evolution of caspase-mediated cell death and differentiation: twins separated at birth. Cell Death Differ 24, 1359–1368, doi:10.1038/cdd.2017.37 (2017).

4 Losick, V. P., Morris, L. X., Fox, D. T. & Spradling, A. Drosophila stem cell niches: a decade of discovery suggests a unified view of stem cell regulation. Dev Cell 21, 159–171, doi:10.1016/j.devcel.2011.06.018 (2011).

5 Tang, H. L., Tang, H. M., Fung, M. C. & Hardwick, J. M. In vivo CaspaseTracker biosensor system for detecting anastasis and non-apoptotic caspase activity. Sci Rep 5, 9015, doi:10.1038/srep09015 (2015).

6 Huang, J. & Kalderon, D. Coupling of Hedgehog and Hippo pathways promotes stem cell maintenance by stimulating proliferation. J Cell Biol 205, 325–338, doi:10.1083/jcb.201309141 (2014).

7 Vied, C. & Kalderon, D. Hedgehog-stimulated stem cells depend on non-canonical activity of the Notch co-activator Mastermind. Development 136, 2177–2186, doi:10.1242/dev.035329 (2009).

8 Zhang, Y. & Kalderon, D. Hedgehog acts as a somatic stem cell factor in the Drosophila ovary. Nature 410, 599–604, doi:10.1038/35069099 (2001).

9 Rojas-Rios, P., Guerrero, I. & Gonzalez-Reyes, A. Cytoneme-mediated delivery of hedgehog regulates the expression of bone morphogenetic proteins to maintain germline stem cells in Drosophila. PLoS Biol 10, e1001298, doi:10.1371/journal.pbio.1001298 (2012).

10 Sahai-Hernandez, P. & Nystul, T. G. A dynamic population of stromal cells contributes to the follicle stem cell niche in the Drosophila ovary. Development 140, 4490–4498, doi:10.1242/dev.098558 (2013).

11 Chang, Y. C., Jang, A. C., Lin, C. H. & Montell, D. J. Castor is required for Hedgehog-dependent cell-fate specification and follicle stem cell maintenance in Drosophila oogenesis. Proc Natl Acad Sci U S A 110, E1734–1742, doi:10.1073/pnas.1300725110 (2013).

12 Briscoe, J. & Therond, P. P. The mechanisms of Hedgehog signalling and its roles in development and disease. Nat Rev Mol Cell Biol 14, 416–429, doi:10.1038/nrm3598 (2013).

13 Dai, W., Peterson, A., Kenney, T., Burrous, H. & Montell, D. J. Quantitative microscopy of the Drosophila ovary shows multiple niche signals specify progenitor cell fate. Nat Commun 8, 1244, doi:10.1038/s41467-017-01322-9 (2017).

14 Singh, T., Lee, E. H., Hartman, T. R., Ruiz-Whalen, D. M. & O’Reilly, A. M. Opposing Action of Hedgehog and Insulin Signaling Balances Proliferation and Autophagy to Determine Follicle Stem Cell Lifespan. Dev Cell 46, 720–734 e726, doi:10.1016/j.devcel.2018.08.008 (2018).

15 Hartman, T. R., Strochlic, T. L., Ji, Y., Zinshteyn, D. & O’Reilly, A. M. Diet controls Drosophila follicle stem cell proliferation via Hedgehog sequestration and release. J Cell Biol 201, 741–757, doi:10.1083/jcb.201212094 (2013).

16 Rosales-Nieves, A. E. & Gonzalez-Reyes, A. Genetics and mechanisms of ovarian cancer: parallels between Drosophila and humans. Semin Cell Dev Biol 28, 104–109, doi:10.1016/j.semcdb.2014.03.031 (2014).

17 Szkandera, J., Kiesslich, T., Haybaeck, J., Gerger, A. & Pichler, M. Hedgehog signaling pathway in ovarian cancer. Int J Mol Sci 14, 1179–1196, doi:10.3390/ijms14011179 (2013).

18 Zeng, C., Chen, T., Zhang, Y. & Chen, Q. Hedgehog signaling pathway regulates ovarian cancer invasion and migration via adhesion molecule CD24✓Cancer 8, 786–792, doi:10.7150/jca.17712 (2017).

19 Ray, A., Meng, E., Reed, E., Shevde, L. A. & Rocconi, R. P. Hedgehog signaling pathway regulates the growth of ovarian cancer spheroid forming cells. Int J Oncol 39, 797–804, doi:10.3892/ijo.2011.1093 (2011).

20 Baena-Lopez, L. A. et al. Novel initiator caspase reporters uncover previously unknown features of caspase-activating cells. Development 145, doi:10.1242/dev.170811 (2018).

21 Arthurton, L, Nahotko, D., Alonso, J. & Baena-Lopez, L. A. Non-apoptotic caspase-dependent regulation of enteroblast quiescence in Drosophila. bioRxiv, doi:10.1101/707380 (2019).

22 Shinoda, N., Hanawa, N., Chihara, T., Koto, A. & Miura, M. Droneindependent basal executioner caspase activity sustains Drosophila imaginal tissue growth. Proc Natl Acad Sci U S A 116, 20539–20544, doi:10.1073/pnas.1904647116 (2019).

23 Laws, K. M. & Drummond-Barbosa, D. Genetic Mosaic Analysis of Stem Cell Lineages in the Drosophila Ovary. Methods Mol Biol 1328, 57–72, doi:10.1007/978-1-4939-2851-4_4 (2015).

24 Baena-Lopez, L. A., Alexandre, C., Mitchell, A., Pasakarnis, L. & Vincent, J. P. Accelerated homologous recombination and subsequent genome modification in Drosophila. Development 140, 4818–4825, doi:10.1242/dev.100933 (2013).

25 Riabinina, O. & Potter, C. J. The Q-System: A Versatile Expression System for Drosophila. Methods Mol Biol 1478, 53–78, doi:10.1007/978-1-4939-6371-3_3 (2016).

26 Hartman, T. R. et al. Novel tools for genetic manipulation of follicle stem cells in the Drosophila ovary reveal an integrin-dependent transition from quiescence to proliferation. Genetics 199, 935–957, doi:10.1534/genetics.114.173617 (2015).

27 Zielke, N. et al. Fly-FUCCI: A versatile tool for studying cell proliferation in complex tissues. Cell Rep 7, 588–598, doi:10.1016/j.celrep.2014.03.020 (2014).

28 Napoletano, F. et al. p53-dependent programmed necrosis controls germ cell homeostasis during spermatogenesis. PLoS Genet 13, e1007024, doi:10.1371/journal.pgen.1007024 (2017).

29 Ouyang, Y. et al. Dronc caspase exerts a non-apoptotic function to restrain phospho-Numb-induced ectopic neuroblast formation in Drosophila. Development 138, 2185–2196, doi:10.1242/dev.058347 (2011).

30 Muro, L, Monser, K. & Clem, R. J. Mechanism of Dronc activation in Drosophila cells. J Cell Sci 117, 5035–5041, doi:10.1242/jcs.01376 (2004).

31 Chai, J. et al. Molecular mechanism of Reaper-Grim-Hid-mediated suppression of DIAP1-dependent Dronc ubiquitination. Nat Struct Biol 10, 892–898, doi:10.1038/nsb989 (2003).

32 Lee, T. V. et al. Drosophila IAP1-mediated ubiquitylation controls activation of the initiator caspase DRONC independent of protein degradation. PLoS Genet 7, e1002261, doi:10.1371/journal.pgen.1002261 (2011).

33 Leulier, F. et al. Systematic in vivo RNAi analysis of putative components of the Drosophila cell death machinery. Cell Death Differ 13, 1663–1674, doi:10.1038/sj.cdd.4401868 (2006).

34 Xu, D. et al. The effector caspases drICE and dcp-1 have partially overlapping functions in the apoptotic pathway in Drosophila. Cell Death Differ 13, 1697–1706, doi:10.1038/sj.cdd.4401920 (2006).

35 Hay, B. A., Wolff, T. & Rubin, G. M. Expression of baculovirus P35 prevents cell death in Drosophila. Development 120, 2121–2129. (1994).

36 Huang, J., Reilein, A. & Kalderon, D. Yorkie and Hedgehog independently restrict BMP production in escort cells to permit germline differentiation in the Drosophila ovary. Development 144, 2584–2594, doi:10.1242/dev.147702 (2017).

37 Lu, X., Liu, S. & Kornberg, T. B. The C-terminal tail of the Hedgehog receptor Patched regulates both localization and turnover. Genes Dev 20, 2539–2551, doi:10.1101/gad.1461306 (2006).

38 Johnson, R. L., Milenkovic, L. & Scott, M. P. In vivo functions of the patched protein: requirement of the C terminus for target gene inactivation but not Hedgehog sequestration. Mol Cell 6, 467–478. (2000).

39 Motzny, C. K. & Holmgren, R. The Drosophila cubitus interruptus protein and its role in the wingless and hedgehog signal transduction pathways. Mech Dev 52, 137–150 (1995).

40 Shyamala, B. V. & Bhat, K. M. A positive role for patched-smoothened signaling in promoting cell proliferation during normal head development in Drosophila. Development 129, 1839–1847 (2002).

41 Buszczak, M. et al. The carnegie protein trap library: a versatile tool for Drosophila developmental studies. Genetics 175, 1505–1531, doi:10.1534/genetics.106.065961 (2007).

42 Jimenez-Sanchez, M. et al. The Hedgehog signalling pathway regulates autophagy. Nat Commun 3, 1200, doi:10.1038/ncomms2212 (2012).

43 Nezis, I. P. et al. Ref(2)P, the Drosophila melanogaster homologue of mammalian p62, is required for the formation of protein aggregates in adult brain. J Cell Biol 180, 1065–1071, doi:10.1083/jcb.200711108 (2008).

44 Bjorkoy, G. et al. Monitoring autophagic degradation of p62/SQSTM1. Methods Enzymol 452, 181–197, doi:10.1016/S0076-6879(08)03612-4 (2009).

45 Rojas-Rios, P. et al. Translational Control of Autophagy by Orb in the Drosophila Germline. Dev Cell 35, 622–631, doi:10.1016/j.devcel.2015.11.003 (2015).

46 Li, Y., Wang, S., Ni, H. M., Huang, H. & Ding, W. X. Autophagy in alcohol-induced multiorgan injury: mechanisms and potential therapeutic targets. Biomed Res Int 2014, 498491, doi:10.1155/2014/498491 (2014).

47 Mauvezin, C. & Neufeld, T. P. Bafilomycin A1 disrupts autophagic flux by inhibiting both V-ATPase-dependent acidification and Ca-P60A/SERCA-dependent autophagosome-lysosome fusion. Autophagy 11, 1437–1438, doi:10.1080/15548627.2015.1066957 (2015).

48 Ding, A. X. et al. CasExpress reveals widespread and diverse patterns of cell survival of caspase-3 activation during development in vivo. Elife 5, doi:10.7554/eLife.10936 (2016).

49 Sun, G. et al. A molecular signature for anastasis, recovery from the brink of apoptotic cell death. J Cell Biol 216, 3355–3368, doi:10.1083/jcb.201706134 (2017).

50 Burgon, P. G. & Megeney, L. A. Caspase signaling, a conserved inductive cue for metazoan cell differentiation. Semin Cell Dev Biol, doi:10.1016/j.semcdb.2017.11.009 (2017).

51 M, H. D., Tashakor, A., O’Connell, E. & Fearnhead, H. O. Apoptosome-dependent myotube formation involves activation of caspase-3 in differentiating myoblasts. Cell Death Dis 11, 308, doi:10.1038/s41419-020-2502-4 (2020).

52 Mille, F. et al. The Patched dependence receptor triggers apoptosis through a DRAL-caspase-9 complex. Nat Cell Biol 11, 739–746, doi:10.1038/ncb1880 (2009).

53 Fombonne, J. et al. Patched dependence receptor triggers apoptosis through ubiquitination of caspase-9. Proc Natl Acad Sci USA 109, 10510–10515, doi:10.1073/pnas.1200094109 (2012).

54 Li, S., Li, S., Wang, B. & Jiang, J. Hedgehog reciprocally controls trafficking of Smo and Ptc through the Smurf family of E3 ubiquitin ligases. Sci Signal 11, doi:10.1126/scisignal.aan8660 (2018).

55 Martin, D. N. & Baehrecke, E. H. Caspases function in autophagic programmed cell death in Drosophila. Development 131, 275–284, doi:10.1242/dev.00933 (2004).

56 Daish, T. J., Mills, K. & Kumar, S. Drosophila caspase DRONC is required for specific developmental cell death pathways and stress-induced apoptosis. Dev Cell 7, 909–915, doi:10.1016/j.devcel.2004.09.018 (2004).

57 Roy, S. & Ingham, P. W. Hedgehogs tryst with the cell cycle. / Cell Sci 115, 4393–4397, doi:10.1242/jcs.00158 (2002).

58 Agathocleous, M., Locker, M., Harris, W. A. & Perron, M. A general role of hedgehog in the regulation of proliferation. Cell Cycle 6, 156–159, doi:10.4161/cc.6.2.3745 (2007).

59 Weaver, B. P. et al. Non-Canonical Caspase Activity Antagonizes p38 MAPK Stress-Priming Function to Support Development. Dev Cell 53, 358–369 e356, doi:10.1016/j.devcel.2020.03.015 (2020).

60 Annibaldi, A., Dousse, A., Martin, S., Tazi, J. & Widmann, C. Revisiting G3BP1 as a RasGAP binding protein: sensitization of tumor cells to chemotherapy by the RasGAP 317-326 sequence does not involve G3BP1. PLoS One 6, e29024, doi:10.1371/journal.pone.0029024 (2011).

61 Moreno, E., Basler, K. & Morata, G. Cells compete for decapentaplegic survival factor to prevent apoptosis in Drosophila wing development Nature 416, 755–759 (2002).

62 Huey, R. B., Wakefield, T., Crill, W. D. & Gilchrist, G. W. Within- and between-generation effects of temperature on early fecundity of Drosophila melanogaster. Heredity (Edinb) 74 (Pt 2), 216–223 (1995).

63 Terashima, J. & Bownes, M. Translating available food into the number of eggs laid by Drosophila melanogaster. Genetics 167, 1711–1719, doi:10.1534/genetics.103.024323 (2004).

64 Terashima, J., Takaki, K., Sakurai, S. & Bownes, M. Nutritional status affects 20-hydroxyecdysone concentration and progression of oogenesis in Drosophila melanogaster. J Endocrinol 187, 69–79, doi:10.1677/joe.1.06220 (2005).

65 Pepling, M. E. Hedgehog signaling in follicle development. Biol Reprod 86, 173, doi:10.1095/biolreprod.112.100859 (2012).

66 Zhou, J., Peng, X. & Mei, S. Autophagy in Ovarian Follicular Development and Atresia. Int J Biol Sci 15, 726–737, doi:10.7150/ijbs.30369 (2019).

67 Dick, S. A. & Megeney, L. A. Cell death proteins: an evolutionary role in cellular adaptation before the advent of apoptosis. Bioessays 35, 974–983, doi:10.1002/bies.201300052 (2013).

68 Lee, R. E., Brunette, S., Puente, L. G. & Megeney, L. A. Metacaspase Yca1 is required for clearance of insoluble protein aggregates. Proc Natl Acad Sci USA 107, 13348–13353, doi:10.1073/pnas.1006610107 (2010).

