## Supplemental Information for "Non-apoptotic caspase activation sustains ovarian somatic stem cell functions by modulating Hedgehog-signalling and autophagy"

**SUPPLEMENTARY INFORMATION**

**Non-apoptotic caspase activity sustains proliferation and differentiation of ovarian somatic cells by modulating Hedgehog-signalling and autophagy.**

**Alessia Galasso^1^, Daria Iakovleva^1^, Luis Alberto Baena-Lopez^1^***

* Author for correspondence

1: Sir William Dunn School of Pathology. University of Oxford. South Parks Road. Oxfordshire, UK. OX13RE

**GENOTYPE DESCRIPTION OF SUPPLEMENTAL FIGURES**

**Supplementary Figure 1.**

**b.** w;; *Dronc*^KO-Gal4^ / UAS-Histone-RFP (BL56555);

**c.** w; UAS-*flippase (BL4539)*/+; *Dronc*^KO-Gal4^ / Actin5C FRT-*stop*-FRT *lacZ*-nls/+ (BL6355)

**d.** *Dronc::*V5::TurboID/+

**e.** *Drice::*V5::TurboID/+

**f-g.** *yw hs*-*flippase*^1.22^/+; FRT80 *Dronc*^I29^/ FRT80 Ubi*GFP*

**Supplementary Figure 2.**

**b.** *109-30*-Gal4 (BL7023)/+; UAS-*Histone-RFP* (BL56555) Tub-*G80^ts^* (BL7019)/+;

**c.** CTRL: *109-30*-Gal4 (BL7023)/QUAS-*CD8-GFP* (BL 30002); *Dronc*^KO^ Tub-G80^ts^ (BL7019) / TM6b.

**d**. *Dronc* -/-: *109-30*-Gal4 (BL7023)/QUAS-*CD8-GFP* (BL 30002); *Dronc*^KO^ Tub-*G80^ts^* (BL7019) / UAS-*flipasse* (BL8209) *Dronc*^KO-FRT-Dronc-GFP-APEX-FRT-QF^

**f.** *ptc-*Gal4(BL2017)/+; UAS-*Histone-RFP* (BL56555) Tub-*G80^ts^* (BL7019)/+;

**g.** CTRL: *ptc-*Gal4 (BL2017) / QUAS-*CD8-GFP* (BL30002); Tub-*G80^ts^* (BL7019) /+.

**h.** *Dronc*-/-: *ptc-*Gal4(BL2017)/QUAS-*CD8-GFP* (BL 30002); *Dronc*^KO^ Tub-*G80^ts^* (BL7019) / UAS-*Flipasse* (BL8209) *Dronc*^KO-FRT-Dronc-GFP-APEX-FRT-QF^

**i.** From left to right: 1: *ptc-*Gal4(BL2017)/+; 2: *ptc-*Gal4 (BL2017)/+; *Dronc*^KO^ Tub-*G80^ts^* (BL7019) /+; 3: *ptc-*Gal4(BL2017)/ UAS-*Dronc* (BL56198); *Dronc*^KO^ Tub-*G80^ts^* (BL7019) /+

**Supplementary Figure 3.**

**a.** *109-30*-Gal4 (BL7023)/+.

**b.** *109-30*-Gal4 (BL7023)/+; UAS-*Ci*-RNAi (BL28984)/+.

**c, d.** From left to right:

CTRL= *109-30*-Gal4 (BL7023)/+

*Dronc* -/- =*109-30*-Gal4 (BL7023)/+; UAS-*Ci*-RNAi (BL28984)/+.

CTRL = *ptc-*Gal4(BL2017)/ + ; Tub-*G80^ts^* (BL7019)/+

UAS-*ptc*^1130X^YFP = *ptc-*Gal4(BL2017)/ UAS-*ptc*^1130X^YFP (BL52215); Tub-*G80^ts^* (BL7019)/+

**e.** *109-30*-Gal4 (BL7023)/UAS-*Ci* (BL32571); *Dronc*^KO^ Tub-*G80^ts^* (BL7019) / +

**f.** *109-30*Gal4 (BL7023)/UAS-*Ci* (BL32571); *Dronc*^KO^ Tub-*G80^ts^* (BL7019)/ UAS-*Flipasse* (BL8209) *Dronc*^KO-FRT-Dronc-GFP-APEX-FRT-QF^

**Supplementary Figure 4.**

**a.** *ptc-*GFP^CB02030^ / +; *Dronc*^KO^ /+

**b.** From left to right: **(***Dronc*^KO^ Tub-*G80^ts^* (BL7019)/+); (*ptc-*Gal4(BL2017)/+); (*ptc-*Gal4(BL2017)/+; *Dronc*^KO^ Tub-*G80^ts^* (BL7019) /+)

**c.** *ptc-*Gal4(BL2017)/ UAS-*Dronc*-RNAi (a gift from Pascal Meier)

**d.** *ptc*^S2^ (BL6332)/+; *Dronc* +/-

**e.** From left to right: **(***Dronc*^KO^ Tub-*G80^ts^* (BL7019)/+); (*ptc-*Gal4(BL2017)/+);(*ptc-*Gal4(BL2017)/+; *Dronc*^KO^ Tub-*G80^ts^* (BL7019) /+)

**f.** *ptc-*Gal4(BL2017)/ UAS-*ptc*^1130X^YFP (BL52215); Tub-*G80^ts^* (BL7019)

**g.** *ptc-*Gal4(BL2017)/ UAS-*ptc*^1130X^YFP (BL52215); *Dronc*^KO^ Tub-*G80^ts^* (BL7019) /+

**h.** From left to right**:**

CTRL = *ptc-*Gal4(BL2017)/ UAS-*ptc*^1130X^ YFP (BL52215); Tub-*G80^ts^* (BL7019)

*Dronc +/-* = *ptc-*Gal4(BL2017)/ UAS-*ptc*^1130X^YFP (BL52215); *Dronc*^KO^ Tub-*G80^ts^* (BL7019) /+

**i.** *ptc-*Gal4(BL2017)/ *ptc-*GFP^CB02030^; *Dronc*^KO^ Tub-*G80^ts^* (BL7019) / +

**SUPPLEMENTARY FIGURE LEGENDS**

**Supplementary Figure 1. Tools and resources used to uncover non-apoptotic caspase activation on the germarium.**

**a.** Design of the temporal caspase reporter Drice-based sensor (DBS-S-QF). Left**:** schematic representation of the membrane attached mCD8-DBS-QF sensor. Right: different labelling systems used in combination with mCD8-DBS-QF to visualize the temporal patterns of apical caspase activation. The current/ongoing activation of caspases is obtained upon induction of QUAS-Tomato-HA. QF translocation can also induce the expression of a Gal4 transcription factor (QUAS-Gal4). Gal4 production can then trigger the expression of a second cellular marker (UAS-CD8-GFP). The appearance of the GFP signal is delayed in time respect to the QUAS-Tomato since it demands a second round of transcriptional events. If apical caspase activation terminates QF translocation finishes; however, the transcriptional amplification obtained with the Gal/UAS transcription loop can maintain the GFP signal for longer after the QUAS-tomato signal disappears (Old caspase activation, green cells). Lineage-tracing (permanent labelling) of caspase activating cells is also achievable by using DBS-QF. In this case the QF transcription factor activates a recombinase (QUAS-*flippase*), which subsequently mediates the genomic excision of a FRT-stop cassette. Before recombination, the FRT-stop cassette prevents the activation of a cellular marker (nuclear-lacZ) under the regulation of an *actin* promoter (permanent labelling).

**b.** Representative confocal image showing the expression pattern of *Dronc*^KO^-Gal4 UAS-Histone-RFP in the germarium (red and gray) at 29°C 10 days after adult eclosion. Dapi stains the nuclei (blue). Notice the expression of *Dronc* in somatic cells of the germarium (arrows) as well as germarium the stalk and polar cells (arrowheads). Scale bars represent 10 µm in the entire figure.

**c.** Representative cell-lineage tracing of cells transcribing *Dronc* in the germarium (green) at 29°C 10 days after adult eclosion. The somatic cells are labelled with FasIII (red). Dapi stains the nuclei (blue). Notice the expression of Dronc in the follicular cells (white arrow), stalk cells (white arrowhead). Experimental flies were kept for 10 days at 29°C after eclosion and prior dissection.

**d**. Biotinylation signal (green) generated in the germarium by a *Dronc*-TurboID allele; notice the signal enrichment in stalk cells (white arrows) and polar cells (white arrowhead). FasIII staining (red) labels the somatic cells and Dapi (blue) the nuclei. Experimental flies were kept for 10 days at 29°C after eclosion and prior dissection.

**e.** Expression pattern of Drice-TurboID protein detected through biotin immunolabelling with Streptavidin (green). Notice the biotinylation enrichment in the somatic cells (including the presumptive follicular stem cells; white arrow), as opposed to the low levels in the germline (white symbol). The somatic cells are labelled with FasIII (red). Dapi labels the DNA (blue). Experimental flies were kept for 10 days at 29°C after eclosion and prior dissection.

**f-g.** Expression of Castor (red) and FasIII (gray) in mutant morphogenetic mosaics for *Dronc^l^*^29^ in the germarium (GFP - cells). Notice the downregulation of Castor (red, white arrows) in g.

**h.** Diagrams showing the conditional *Dronc* alleles used throughout the manuscript.

**Supplementary Figure 2. Expression pattern of the relevant Gal4 drivers.**

**a.** Schematic summarising the presumptive cells activating the *109-30*-Gal4 driver (coloured in red) in the germarium at 29°C.

**b.** Representative confocal image expressing UAS-Histone-RFP (red) under the regulation of *109-30*-Gal4 driver; Castor (green) and Dapi (blue) stainings label the follicular cells and the nuclei, respectively. Scale bars represent 10 µm in the entire figure. In the entire figure, experimental flies were kept for 10 days at 29°C after eclosion and prior dissection.

**c.** Wildtype expression of the follicular marker Castor (red and/or gray) in a representative germarium of the following genotype *109-30-Gal4/+; UAS-Histone-RFP Dronc^KO^ Tub-G80^ts^/+.*

**d.** Castor expression (red, gray and white arrows) at 29°C in a *Dronc* mutant germarium of the following genotype *109-30-Gal4/QUAS-CD8-GFP; Dronc^KO^ Tub-G80^ts^/ UAS-flippase Dronc^KO-FRT-Dronc-GFP-APEX-FRT-QF^*. *Dronc*-expressing cells excising the rescue cassette are labelled with GFP (green); notice the reduction in number of Castor-expressing cells in the follicular cell domain (white arrows).

**e.** Schematic depicting the presumptive cells activating the *ptc*-Gal4 driver (coloured in red) in the germarium at 29°C.

**f.** Representative confocal image expressing UAS-Histone-RFP (red) under the regulation of *ptc*-Gal4 driver after 10 days at 29 ^o^C; Castor (green) and Dapi (blue) stainings label the follicular cells and the nuclei, respectively. Scale bars represent 10 µm in the entire figure.

**g.** Wildtype expression of Castor (red and/or gray) in a representative germarium of the following genotype *ptc-Gal4/+; UAS-Histone-RFP Dronc^KO^ Tub-G80^ts^/+.*

**h.** Castor expression (red, gray and white arrows) in a representative *Dronc* mutant germarium of the following genotype *ptc-Gal4/QUAS-CD8-GFP; Dronc^KO^ Tub-G80^ts^/ UAS-flippase Dronc^KO-FRT-Dronc-GFP-APEX-FRT-QF^*. *Dronc*-expressing cells excising the rescue cassette are labelled with GFP (green); notice the reduction in number of Castor-expressing cells (white arrows).

**i.** Percentage of germaria showing TUNEL positive cells in the FasIII domain of the following genotypes *ptc-*Gal4/+; *Tub-G80^ts^* /+ (n=23); *ptc*-Gal4/+; *Dronc^KO^ Tub-G80^ts^*/ *UAS-flippase Dronc^KO-FRT-Dronc-GFP-APEX-FRT-QF^* (n=20); *ptc*-Gal4/UAS-*Dronc*/+; *Tub-G80^ts^* /+ (n=26).

**Supplementary Figure 3. Hh-signaling sustains follicular cell proliferation and differentiation**

**a.** Confocal image showing FasIII (green and/or gray), Castor (red and/or gray), and Ci-155 (blue and/or gray) expression in a control germarium (*109-30-Gal4/+; Tub-G80^ts^/+*). Notice the disruption in Castor expression and the reduction of the FasIII domain (white arrow). Scale bars represents 10 µm in all of the confocal images of the figure. In the entire figure, experimental flies were kept for 10 days at 29°C after eclosion and prior dissection.

**b.** Confocal image showing FasIII (green and/or gray), Castor (red and/or gray), and Ci-155 (blue and/or gray) expression in a representative control germarium (*109-30-Gal4/* UAS*-Ci-*RNAi*; Tub-G80^ts^/+* ).

**c.** Quantification of total number of follicular cells (left) or Castor-expressing cells (right) within the FasIII cellular domain in the following genotypes from left to right: *109-30*-Gal4/+; *Tub-G80*^ts^ (n=15); *109-30*-Gal4/UAS-*Ci*-RNAi; *Tub-G80*^ts^ (n=21); *ptc*-Gal4/+; *Tub-G80^ts^* /+ (n=19); *ptc*-Gal4/UAS-*Ptc^1130X^-YFP*; *Tub-G80^ts^ /* + (n=8). Statistical significance was established by using ordinary Unpaired T-test (****p≤0.0001; ***p≤0.001). Median and quartiles are shown in the violin plot.

**d.** Percentage of Castor-expressing cells versus the total number of Follicular cells in germaria of the genotypes indicated in c (FasIII^+^ cells). Data are expressed as box-and-whiskers plots, with min to max range as whiskers. Statistical significance was established by using using unpaired parametric (plot 1-2) and non-parametric (plots 3-4) T-test (****p≤0.0001). *n* number is shown in c.

**e-f.** Castor expression (red and/or gray) in a representative mutant germarium of the following genotypes: *109-30-Gal4/UAS-Ci; Dronc^KO^ Tub-G80^ts^/ +* (D) *and 109-30-Gal4/UAS-Ci; Dronc^KO^ Tub-G80^ts^/ UAS-flippase Dronc^KO-FRT-Dronc-GFP-APEX-FRT-QF^* (E). Dapi staining labels the nuclei in blue.

**Supplementary Figure 4. Caspase deficiency compromises Hh-signaling activation by inducing the accumulation of Ptc.**

**a.** Representative confocal image showing the expression of Ci-155 (blue and/or gray), *ptc*-GFP (green and/or gray) and Castor (red) in a double heterozygous *ptc-Dronc* mutant germaria of the following genotype *ptc*-*GFP*/+; *Dronc^KO^ Tub-G80^ts^* /+. Scale bars represents 10 µm in all of the confocal images of the figure. In the entire figure, experimental flies were kept for 10 days at 29°C after eclosion and prior dissection.

**b.** Western blot showing the protein levels of Ci-155 (upper lane) and Actin (bottom lane) in germaria of the following genotypes: *Dronc*^KO^/+; *ptc*-Gal4/+; *ptc*-Gal4/+; *Dronc*^KO^/+.

**c-d.** Castor expression (red and/or gray) in representative mutant germaria of the following genotypes: *ptc-Gal4/*UAS*-Dronc-*RNAi*; Dronc^KO^ Tub-G80^ts^/ +* (C) *and ptc^S2^/+; Dronc^KO^ Tub-G80^ts^/+* (E). Dapi labels the DNA (blue).

**e.** Estimation of Ptc-positive punctae size in germaria of of the following genotypes: *Dronc*^KO^/+ (n=10 ); *ptc*-Gal4/+ (n= 9 ); *ptc*-Gal4/+; *Dronc*^KO^/+ (n= 10 ).; the statistical significance between groups was established using one way ANOVA Tukey's multiple comparisons test (****p≤0.0001, ***p≤0.001, *p≤0.05). Notice the enlargement of Ptc-positive particles in a double heterozygous germaria (*ptc*-Gal4/+; *Dronc* +/-). The average value and the standard deviation S.D are indicated by the red lines.

**f-g.** Expression of the *ptc*^1130X^YFP (green) in a representative germarium of the following genotypes: (F) *ptc-Gal4/*UAS*-ptc*^1130X^-YFP *; Tub-G80^ts^/ + and ptc-Gal4/*UAS*-ptc*^1130X^-YFP *; Dronc^KO^ Tub-G80^ts^/ +* (G). Dapi (blue) and FasIII (red) stainings label the nuclei and follicular cells, respectively. Notice the accumulation of GFP signal in G.

**h.** Quantification of *ptc*^1130X^-YFP expression levels in germaria of the following genotypes: *ptc-Gal4/*UAS*-ptc*^1130X^-YFP *; Tub-G80^ts^/ +* (n=8) *and ptc-Gal4/*UAS*-ptc*^1130X^-YFP *; Dronc^KO^ Tub-G80^ts^/ +* (n=16); unpaired T-Test was used to establish the statistical significance (**** p≤0.0001). Median and quartiles are shown in the violin plot.

**i.** Representative confocal image showing the expression of Ci-155 (blue and/or gray), *ptc*-GFP (green and/or gray) and Castor (red) in germaria of the following genotype *ptc*-*GFP*/ *ptc*-*Gal4*; *Dronc^KO^ Tub-G80^ts^* /+. Notice that the expression levels of Ci, *ptc*-GFP and Castor are largely restored.

**KEY RESOURCES TABLE**

| REAGENT or RESOURCE | SOURCE | IDENTIFIER |
| --- | --- | --- |
| Antibodies | | |
| rabbit monoclonal anti-HA (clone C29F4) | Cell Signaling | #3724 |
| rabbit anti-Castor | gift from Alex Gould |  |
| DAPI Solution (1 mg/mL) | ThermoFisher Scientific | 62248 |
| mouse monoclonal anti-β-Gal | Promega | Z378B |
| chicken polyclonal anti-β-Gal | Abcam | ab9361 |
| mouse monoclonal anti-FasIII | Hybridoma Bank | 7G10 |
| rat monoclonal anti-Ci-155-full length | Hybridoma Bank | 2A1 |
| mouse monoclonal anti-Ptc | Hybridoma Bank | Apa1 |
| mouse monoclonal anti-actin | Hybridoma Bank | JLA20 |
| rabbit polyclonal anti-Ref2P | Abcam | ab178440 |
| mouse monoclonal anti-Caspase-9 (C9) | Cell Signaling | #9508 |
| rabbit polyclonal anti-SQSTM1 / P62 | GeneTex | GTX111393 |
| Donkey anti-Rabbit IgG (H+L) Highly Cross-Adsorbed Secondary Antibody, Alexa Fluor 488 | ThermoFisher Scientific | A-21206 |
| Donkey anti-Rabbit IgG (H+L) Highly Cross-Adsorbed Secondary Antibody, Alexa Fluor 555 | ThermoFisher Scientific | A-31572 |
| Donkey anti-Rabbit IgG (H+L) Highly Cross-Adsorbed Secondary Antibody, Alexa Fluor 647 | ThermoFisher Scientific | A-31573 |
| Donkey anti-Mouse IgG (H+L) Highly Cross-Adsorbed Secondary Antibody, Alexa Fluor 488 | ThermoFisher Scientific | A-21202 |
| Donkey anti-Mouse IgG (H+L) Highly Cross-Adsorbed Secondary Antibody, Alexa Fluor 555 | ThermoFisher Scientific | A-31570 |
| Donkey anti-Mouse IgG (H+L) Highly Cross-Adsorbed Secondary Antibody, Alexa Fluor 647 | ThermoFisher Scientific | A-31571 |
| Goat anti-Rat IgG (H+L) Cross-Adsorbed Secondary Antibody, Alexa Fluor 647 | ThermoFisher Scientific | A-21247 |
| Goat anti-Rat IgG (H+L) Cross-Adsorbed Secondary Antibody, Alexa Fluor 555 | ThermoFisher Scientific | A-21434 |
| Streptavidin, Alexa Fluor™ 488 conjugate | ThermoFisher Scientific | S11223 |
| Streptavidin, Alexa Fluor™ 647 conjugate | ThermoFisher Scientific | S21374 |
| mouse monoclonal anti-β-Actin−Peroxidase | Sigma | A3854 |
| Bacterial and Virus Strains | | |
| *Mix & Go!* *E.coli* Transformation Kit | Zymo Research | T3001 |
| Biological Samples |  |  |
| Chemicals, Peptides, and Recombinant Proteins | | |
| RPMI | Sigma | R8758 |
| Gibco™ Fetal Bovine Serum, qualified, heat inactivated | Fisher Scientific | 11550356 |
| Bafilomycin A1 | Merck Chemicals | 19-148 |
| IGEPAL CA-630 | Sigma | I8896 |
| Halt™ Protease and Phosphatase Inhibitor Cocktail | Thermo Fisher Scientific | 78444 |
| BaseMuncher | Expedeon Ltd | BM0025-EXP |
| Protein Assay Dye Reagent | Bio-Rad | #5000006 |
| NuPAGE™ LDS Sample Buffer | Thermo Fisher Scientific | NP0007 |
| NuPAGE™ MOPS SDS Running Buffer | Thermo Fisher Scientific | NP0001 |
| NuPAGE™ 4 to 12%, Bis-Tris | Thermo Fisher Scientific | NP0321PK2 |
| Trans-Blot Turbo Mini 0.2 µm Nitrocellulose Transfer | Bio-Rad | #1704158 |
| SuperSignal™ West Pico PLUS Chemiluminescent Substrate | Thermo Fisher Scientific | 34580 |
| CL-XPosure™ Film | Thermo Fisher Scientific | 34090 |
| Poly-Prep Slides | Sigma | P0425-72EA |
| Aqua-Poly/Mount | Polysciences, Inc | 18606 |
| Critical Commercial Assays | | |
| DeadEnd™ Colorimetric TUNEL System | Promega | G7361 |
| Click-iT™ Plus EdU Cell Proliferation Kit for Imaging, Alexa Fluor™ 647 dye | Thermo Fisher Scientific | C10640 |
| RNeasy Plus kit | Qiagen | 74034 |
| Maxima First Strand cDNA synthesis | ThermoFisher Scientific | K1642 |
| QuantiNova SYBR Green | Qiagen | 208054 |
| Deposited Data | | |
| Experimental Models: Cell Lines | | |
| OVCAR-3 | ATCC (gift from David Carter) | NIH:OVCAR-3 [OVCAR3] (ATCC^®^ HTB-161^™^) |
| Experimental Models: Organisms/Strains | | |
| w[1118]; P{w[+mC]=QUAS-FLPo.P}1 | Bloomington Drosophila Stock Center | 30126 |
| P{ry[+t7.2]=Act5C(FRT.polyA)lacZ.nls1}3, ry[506] | Bloomington Drosophila Stock Center | 6355 |
| w[*]; P{w[+mC]=wor.GAL4.A}2, P{w[+mC]=UAS-mira.GFP}1.2/CyO, P{w[+mC]=ActGFP}JMR1; P{w[+mC]=UAS-His-RFP}3 | Bloomington Drosophila Stock Center | 56555 |
| *Dronc*^TurboID^ | Shinoda N. et al., 2019 |  |
| *Drice*^TurboID^ | Shinoda N. et al., 2019 |  |
| y[1] w[*]; P{w[+mW.hs]=GawB}109-30/CyO | Bloomington Drosophila Stock Center | 7023 |
| w[*]; P{w[+mC]=tubP-GAL80[ts]}20; TM2/TM6B, Tb[1] | Bloomington Drosophila Stock Center | 7019 |
| y[1] w[1118]; P{w[+mC]=UAS-FLP.Exel}3 | Bloomington Drosophila Stock Center | 8209 |
| w[*]; P{w[+mW.hs]=GawB}ptc[559.1] | Bloomington Drosophila Stock Center | 2017 |
| w[1118]; P{w[+mC]=Ubi-GFP.E2f1.1-230}19 P{w[+mC]=Ubi-mRFP1.NLS.CycB.1-266}15/CyO, P{ry[+t7.2]=en1}wg[en11]; MKRS/TM6B, Tb[+] | Bloomington Drosophila Stock Center | 55123 |
| w[*]; P{w[+mC]=UAS-p35.H}BH1 | Bloomington Drosophila Stock Center | 5072 |
| w[*]; P{w[+mC]=UAS-p35.H}BH2 | Bloomington Drosophila Stock Center | 5073 |
| y[1] w[*]; P{w[+mC]=UAS-FLAG-smo.act}2 | Bloomington Drosophila Stock Center | 44621 |
| y[1] v[1]; P{y[+t7.7] v[+t1.8]=TRiP.JF01715}attP2 | Bloomington Drosophila Stock Center | 28984 |
| y[1] sc[*] v[1] sev[21]; P{y[+t7.7] v[+t1.8]=TRiP.HMC03872}attP40 | Bloomington Drosophila Stock Center | 55686 |
| w[*]; P{w[+mC]=UAS-Dronc.FLAG}2 | Bloomington Drosophila Stock Center | 56198 |
| y[1] sc[*] v[1] sev[21]; P{y[+t7.7] v[+t1.8]=TRiP.GL00047}attP2/TM3, Sb[1] | Bloomington Drosophila Stock Center | 35177 |
|  | Bloomington Drosophila Stock Center |  |
| y[1] w[1118]; P{w[+mC]=QUAS-mCD8-GFP.P}5J | Bloomington Drosophila Stock Center | 30002 |
| w[*]; P{w[+mC]=UAS-ptc.1130X.YFP}2 | Bloomington Drosophila Stock Center | 52215 |
| y[1]; P{ry[+t7.2]=neoFRT}42D ptc[S2]/CyO | Bloomington Drosophila Stock Center | 6332 |
| ptc-GFP^CB02030^ | a gift from Isabel Guerrero |  |
| UAS-DroncRNAi | a gift from Pascal Meier |  |
| UAS-DriceRNAi UAS-DecayRNAi | a gift from Pascal Meier |  |
| UAS-DammRNAi, UAS-Dcp1RNAi | a gift from Pascal Meier |  |
| Dronc^KO-Gal4^ | doi.org/10.1101/707380 |  |
| Dronc^KO-FRT-Dronc-GFP-APEX-FRT-QF^ | doi.org/10.1101/707380 |  |
| Dronc^KO-FRT-Dronc-GFP-APEX-FRT-Suntag-HA-Cherry^ | doi.org/10.1101/707380 |  |
| Dronc^KO-FRT-Dronc-GFP-APEX-FRT-FL- Dronc-CAE- Suntag-HA-Cherry^ | doi.org/10.1101/707380 |  |
| y[1] w[*]; P{w[+mC]=UAS-FLP.D}JD1 |  | 4539 |
| Oligonucleotides | | |
| ON-TARGETplus Human CASP9 siRNA | Horizon | L-003309-00-0005, 842 |
| ON-TARGETplus Human PTCH1 siRNA | Horizon | L-003924-00-0005, 5727 |
| ON-TARGET plus Non-targeting Pool | Horizon | D-001810-10-05 |
| *Patched1* Forward CCACGACAAAGCCGACTACAT | Xiaoyun Liao et al., 2009 |  |
| *Patched1* Reverse GCTGCAGATGGTCCTTACTTTTTC | Xiaoyun Liao et al., 2009 |  |
| β*-actin* Forward CCTGGCACCCAGCACAAT | Xiaoyun Liao et al., 2009 |  |
| β*-actin Reverse* GGGCCGGACTCGTCATAC | Xiaoyun Liao et al., 2009 |  |
| Recombinant DNA | | |
| Software and Algorithms | | |
| ImageJ 1.52n | <http://fiji.sc/> | Version 1.52n |
| Flybases | <https://flybase.org> |  |
| Graphpad Prism | <http://www.graphpad.com/> | Version 8.4.3 |
| Adobe Photoshop 2020 | Adobe |  |
| Adobe Illustrator 2020 | Adobe |  |
| Other | | |
