## Supplemental Figures for "Non-apoptotic caspase activation sustains ovarian somatic stem cell functions by modulating Hedgehog-signalling and autophagy"

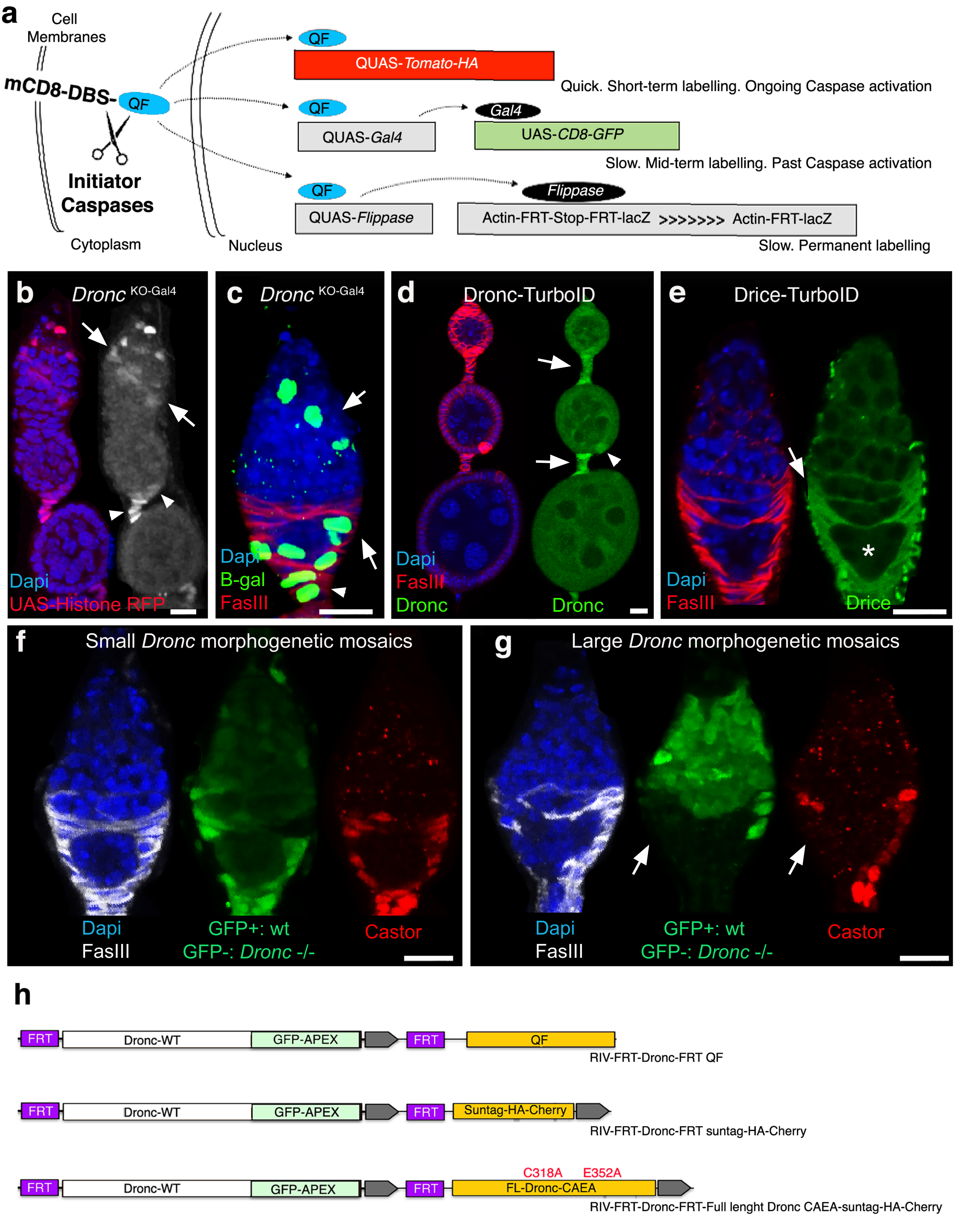


**Galasso et al., supp figure 1**

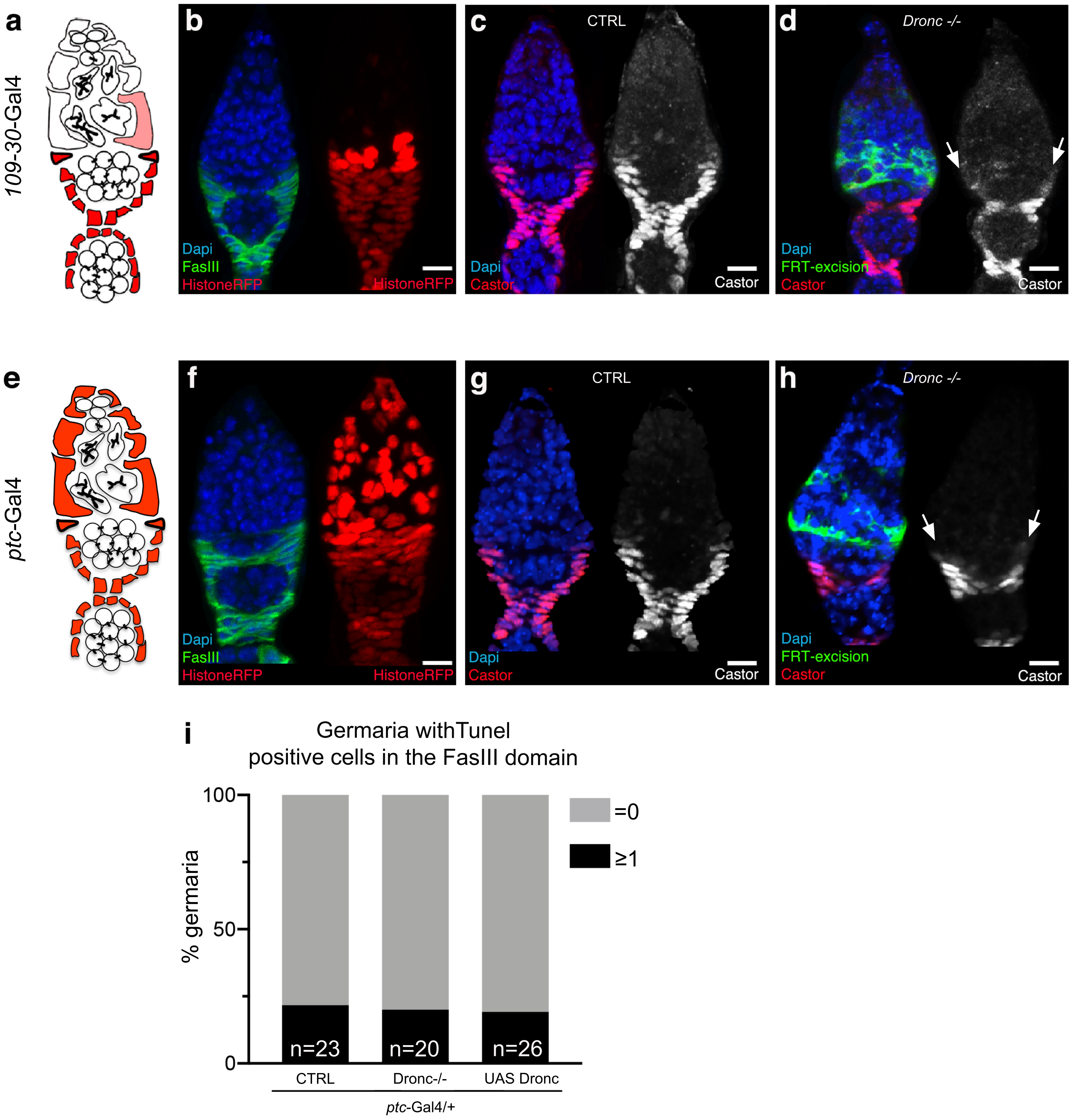


**Galasso et al supp figure 2**

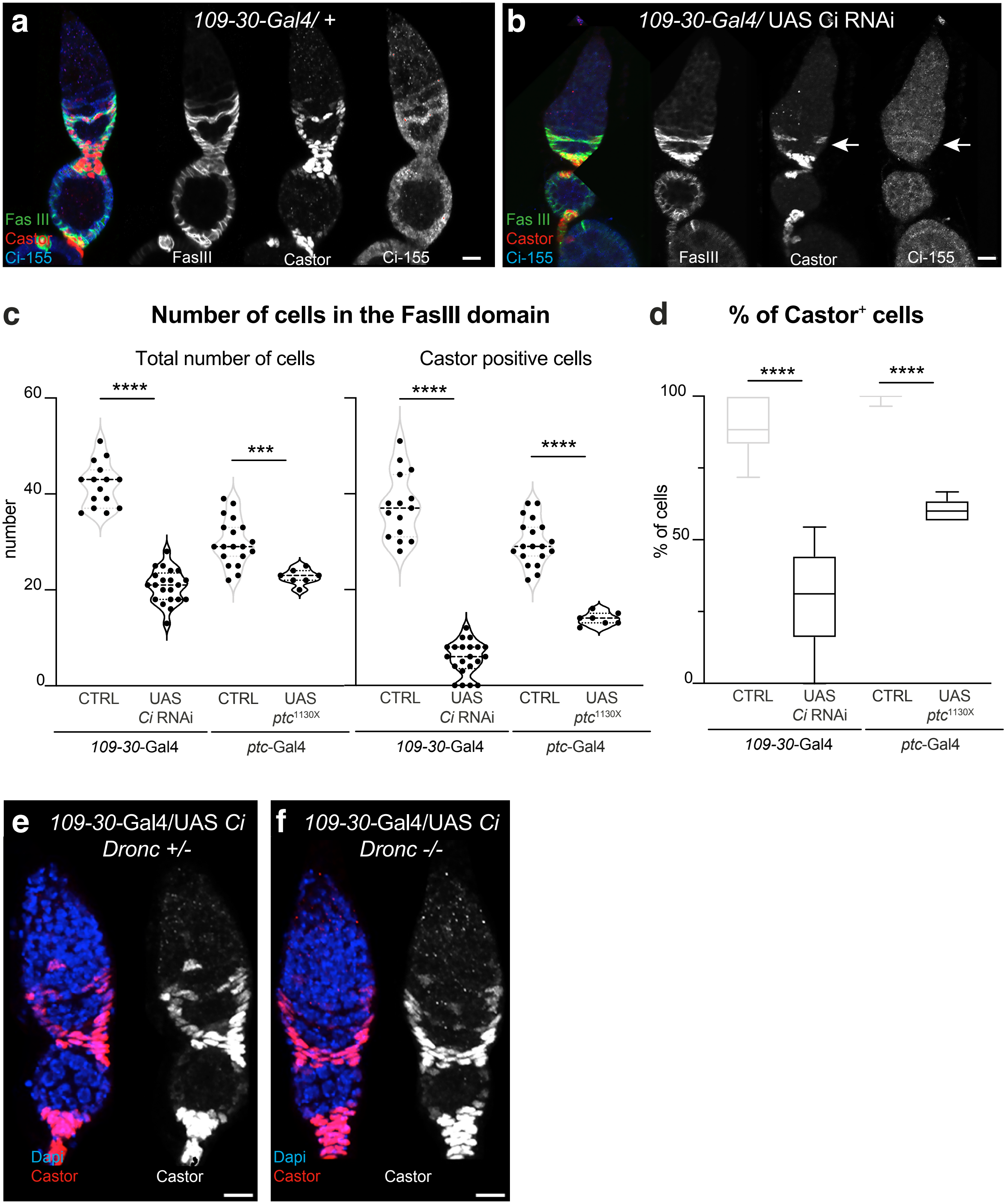


Galasso et al., suppl. fig 3

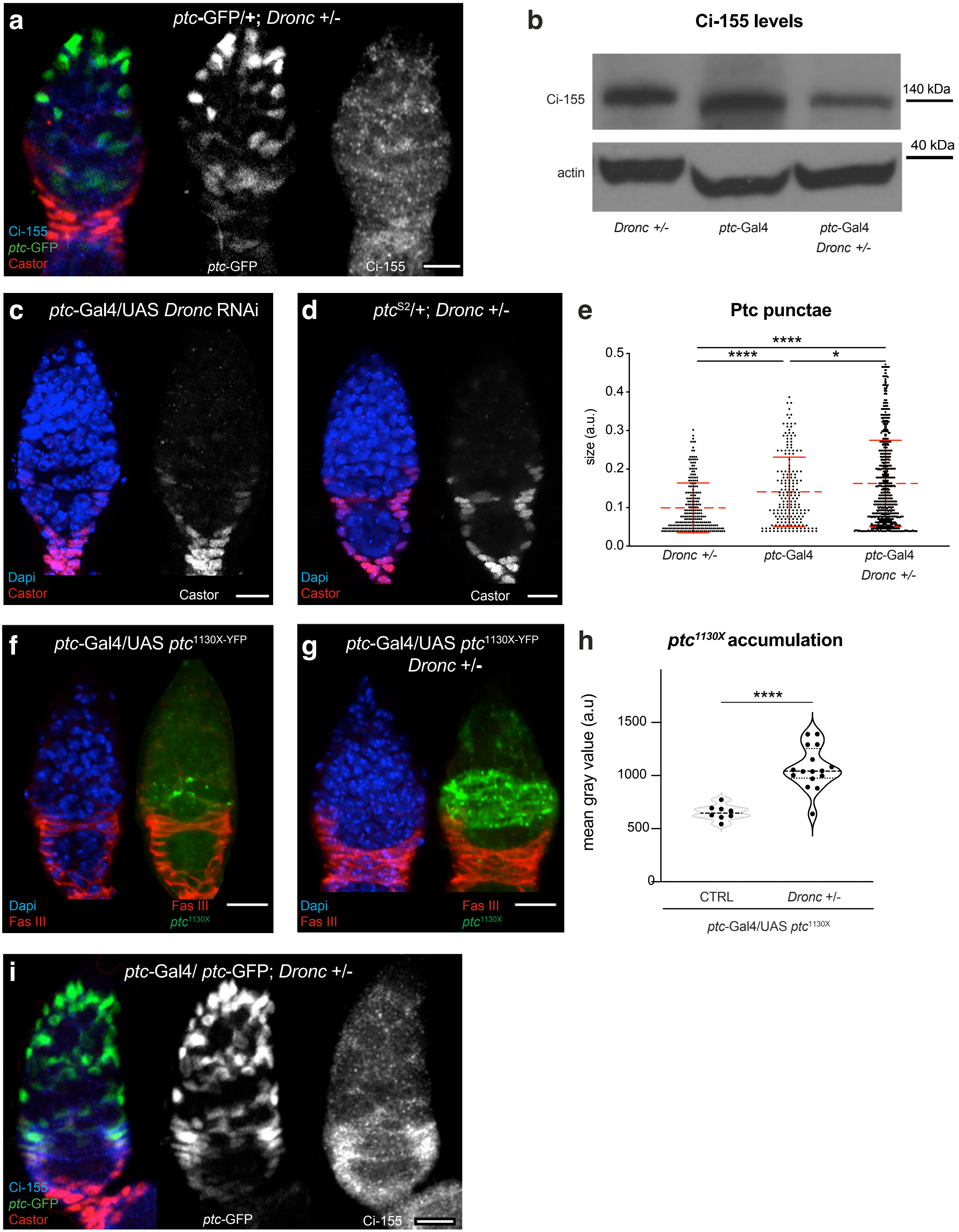


**Galasso et al., suppl. fig 4**
